# Tissular glucocorticoid reactivating enzyme 11β-HSD1 drives pathogenic myofibroblast differentiation in chronic kidney disease

**DOI:** 10.1101/2025.01.09.631985

**Authors:** Grégoire Arnoux, David Legouis, Matthieu Tihy, Raphaël Yvon, Nicolas Liaudet, Seraina O. Moser, Hélène Poinot, Ali Sassi, Olga M. Lempke, Marylise Fernandez, Isabelle Monnay, Alexandra Chassot, Carole Bourquin, Solange Moll, Emmanuel Somm, Joseph Rutkowski, Roland H. Wenger, Maarten Naesens, Alex Odermatt, Stellor Nlandu Khodo, Aurélien Pommier, Eric Feraille

## Abstract

Chronic kidney disease (CKD) is a growing public health crisis, affecting over 10% of the global population and significantly increasing mortality and morbidity. Irrespective of its underlying cause, tubulointerstitial fibrosis (TIF) is a hallmark of CKD progression, with myofibroblasts being the primary effectors of renal fibrosis.

Here, we show that 11beta-hydroxysteroid dehydrogenase type 1 (11β-HSD1) is a critical driver of pathogenic myofibroblast differentiation and fibrosis in CKD. Using genetic deletion and pharmacological inhibition of 11β-HSD1 in mouse models, we demonstrate a marked reduction in TIF severity and improved renal function, linked to the suppression of a regulatory myofibroblast (Reg-MF) subpopulation.

Single-cell and spatial transcriptomics data reveal that 11β-HSD1 is essential for the activation and expansion of Reg-MFs, which is conserved across species and predicts worse outcomes in CKD patients and kidney allograft recipients.

These findings establish a direct link between 11β-HSD1 activity and renal fibrogenesis, highlighting its role during the transition from pericytes to pathogenic Reg-MFs. Our results support 11β-HSD1 inhibition as a promising therapeutic strategy to mitigate CKD progression, offering both mechanistic insights and translational potential for improving patient outcomes.

## Introduction

Chronic kidney disease (CKD) is a critical public health issue^1,2^ affecting 10 to 15% of the global population^3^ and contributing significantly to morbidity and mortality^1^. A key pathological feature of CKD is tubulointerstitial fibrosis (TIF), a process marked by excessive extracellular matrix (ECM) deposition and remodeling of normal tissue architecture^4^, ultimately impairing kidney function^5^. Due to incomplete knowledge of the cellular and molecular drivers of fibrosis, a curative antifibrotic therapy remains elusive. Myofibroblasts are central to ECM production and TIF in CKD. Myofibroblasts are derived from multiple interstitial resident cell types, including resident fibroblasts and pericytes, which together participate in the myofibroblast heterogeneity or “myofibroblast mosaic”^6,7^. These cells contribute differently to CKD progression, suggesting functional specialization within myofibroblast subpopulations. Yet, the precise mechanisms driving their differentiation and pathogenic activity are not fully understood^3,6,8,9^. Among potential regulators, the enzymes 11beta-hydroxysteroid dehydrogenase type 1 and type 2 (11β-HSD1 and 11β-HSD2) play key roles in the modulation of local glucocorticoid bioavailability via the regulation of their intracellular levels in non-adrenal organs^10,11^. While 11β-HSD2 plays a protective role in kidney function by avoiding excessive glucocorticoid-induced mineralocorticoid receptor activity^11^, the role of 11β-HSD1 in CKD is less well understood. Evidence from other organ systems suggests that 11β-HSD1 promotes fibrotic processes, as seen in the skin and myocardium, where its activity has been linked to pathological tissue remodeling^12,13^ and more recently to fibroblast differentiation^14^. In this study, we investigated the role of 11β-HSD1, which regenerates active glucocorticoids, in the pathogenesis of the fibrotic process characterizing CKD. Using genetic deletion and pharmacological inhibition of 11β-HSD1 in two different mouse models of CKD, we investigated its role in TIF, renal function and myofibroblast heterogeneity. Our findings reveal that 11β-HSD1 is essential for the generation of a regulatory myofibroblast (Reg-MF) population that exacerbates fibrosis. We show that inhibition of 11β-HSD1 significantly improves renal structure and function in CKD. By elucidating the mechanism of 11β-HSD1-driven fibrosis and its therapeutic implications, this study provides a new insight for targeting tissular glucocorticoid activation as a novel strategy to mitigate CKD progression.

## Results

### Renal mesenchymal cells exhibit high 11β-HSD1 expression during CKD progression

To investigate the expression and localization of 11β-HSD1 during CKD, we performed immunofluorescence and confocal microscopy on whole kidney transversal sections. In healthy kidneys 11β-HSD1 protein was predominantly localized in the proximal tubules, particularly along the outer medulla (S3 part), as previously described^15^. Additionally, we detected intense labelling of well-organized peritubular interstitial cells mainly located both in the kidney cortex and medulla (Fig. 1a). During chronic kidney injury, we observed a marked redistribution and upregulation of interstitial 11β-HSD1 expression in injured cortical areas, while tubular cell expression of 11β-HSD1 was markedly decreased (Fig. 1b, Extended Data Fig.1a and 1b). 3D immunofluorescence reconstruction imaging and spatial distribution analyses using Ripley’s K-function confirmed a non-random distribution with a shortening distance between 11β-HSD1 positive cells under pathological conditions (Fig. 1c and d, Extended data Fig1.c). To further characterize these cells, we performed high-resolution confocal microscopy with co-staining of 11β-HSD1 and platelet-derived growth factor receptor beta (PDGFR-β), a well-established marker of mesenchymal cells^16^. Virtually all interstitial 11β-HSD1 expressing cells also expressed PDGFR-β, with negligible glomerular expression (Fig. 1e and f, Extended Data Fig.1d-f). We then performed single-nucleus RNA sequencing (snRNAseq) on intact and injured kidneys, identifying all major renal cell types (Fig. 1g, Extended data Fig.1g). Consistent with the immunofluorescence data, *Hsd11b1* expression was found in proximal tubules, while the fibroblastic compartment displayed the highest expression levels of the enzyme (Fig. 1h, Extended Data Fig.1h). Together, these findings highlight a previously underappreciated localization of 11β-HSD1 in mesenchymal cells. The marked upregulation and redistribution of 11β-HSD1/PDGFR-β-positive interstitial cells in CKD suggest a potential role for these cells in TIF and renal injury response.

**Figure 1.**
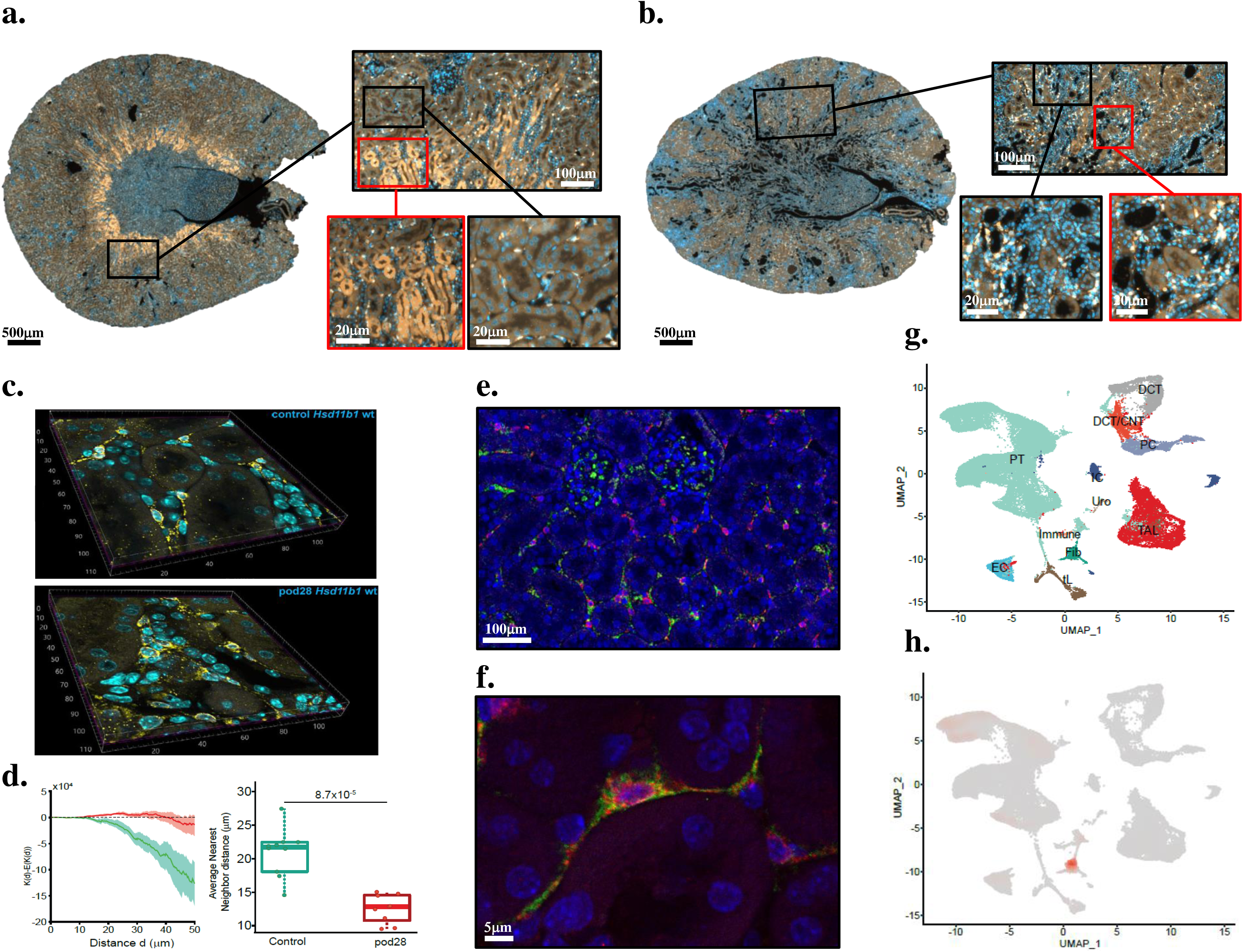
A subpopulation of renal PDGFRβ-positive fibroblasts highly expresses 11β-HSD1. (a and b) Representative immunofluorescence images of 11β-HSD1/DAPI staining of control (**a**) and pod28 post-injury (**b**). 11β-HSD1, gold; DAPI, blue. (**c**) Representative 3D reconstruction from cortical confocal images illustrating cortical interstitial cells expressing 11β-HSD1 in control (upper) and pod (lower) mice kidneys. 11β-HSD1, gold; DAPI, blue. (**d**) 3D spatial analysis of 11β -HSD1-expressing cells. Box plots depicting the average distance to the nearest neighbor (left panel). The distance between the two closest 11β-HSD1-expressing cells is significantly reduced in pod mice compared to control mice (right panel). (**e and f**) Representative confocal images of 11β-HSD1/PDGFRβ/DAPI in control mice kidney sections. 11β-HSD1, red; PDGFRβ, green; DAPI, blue. (**g**) UMAP projection of the integrated snRNAseq dataset, including wt and *Hsd11b1* ko animals, with and without injury (pod), identifying major renal cell types. Eleven mice were included. PT, proximal tubule; tL and TAL, thin and thick ascending limb; DCT, distal convoluted tubule; DCT/CNT, distal convoluted/connecting tubules; PC, principal cell; IC, intercalated cells; Fib, fibroblast; Immune, immune cell; Uro, urothelial cell; EC, endothelial cell; UMAP, uniform manifold approximation and projection. (**h**) Gene-weighted density expression of *Hsd11b1* embedded in the UMAP plot.

### Genetic ablation of 11β-HSD1 improves renal structure and function in CKD

According to the pro-fibrotic role of 11β-HSD1 in various organs^12,17^, we investigated its impact in CKD progression. We generated constitutive *Hsd11b1* knockout (ko) mice and confirmed 11β-HSD1 ablation in the kidney on the mRNA, protein and functional levels (Extended Data Fig.2a-d). snRNAseq analysis revealed reduced activity of the cellular corticosteroid signaling pathway across all kidney cell types in ko compared to the wild-type (wt) mice (Extended Data Fig.2e). Baseline renal parameters, including glomerular filtration rate (GFR), kidney size, morphology and blood pressure, were comparable between the *Hsd11b1* wt and ko animals (Extended data Fig.2f and g). We then took advantage of the POD-ATTAC mouse model (pod), which generates glomerular injury via targeted and inducible podocyte ablation^14^. (Fig. 2a). Genetic ablation of 11β-HSD1 ameliorated the CKD phenotype assessed 4 weeks after injury, as shown by higher GFR (Fig. 2b) and markedly reduced renal TIF, regardless of the severity of tubular or glomerular injury (Fig.2c-h and Extended data Fig.2h-i). Similarly, in the unilateral ureteral obstruction (UUO) model, which causes tubular injury and subsequent fibrosis through increased intratubular hydrostatic pressure^8^, *Hsd11b1* ko animals displayed significantly attenuated fibrosis compared to wt controls 7 days after ureteral ligation (Extended Fig.2j-m). In both models, automated quantification of immunofluorescence imaging showed a substantial reduction in PDGFR-β-positive fibroblasts in the cortical interstitial regions of injured ko kidneys compared to wt animals (Fig.2i-j, Extended data Fig.2n).

**Figure 2.**
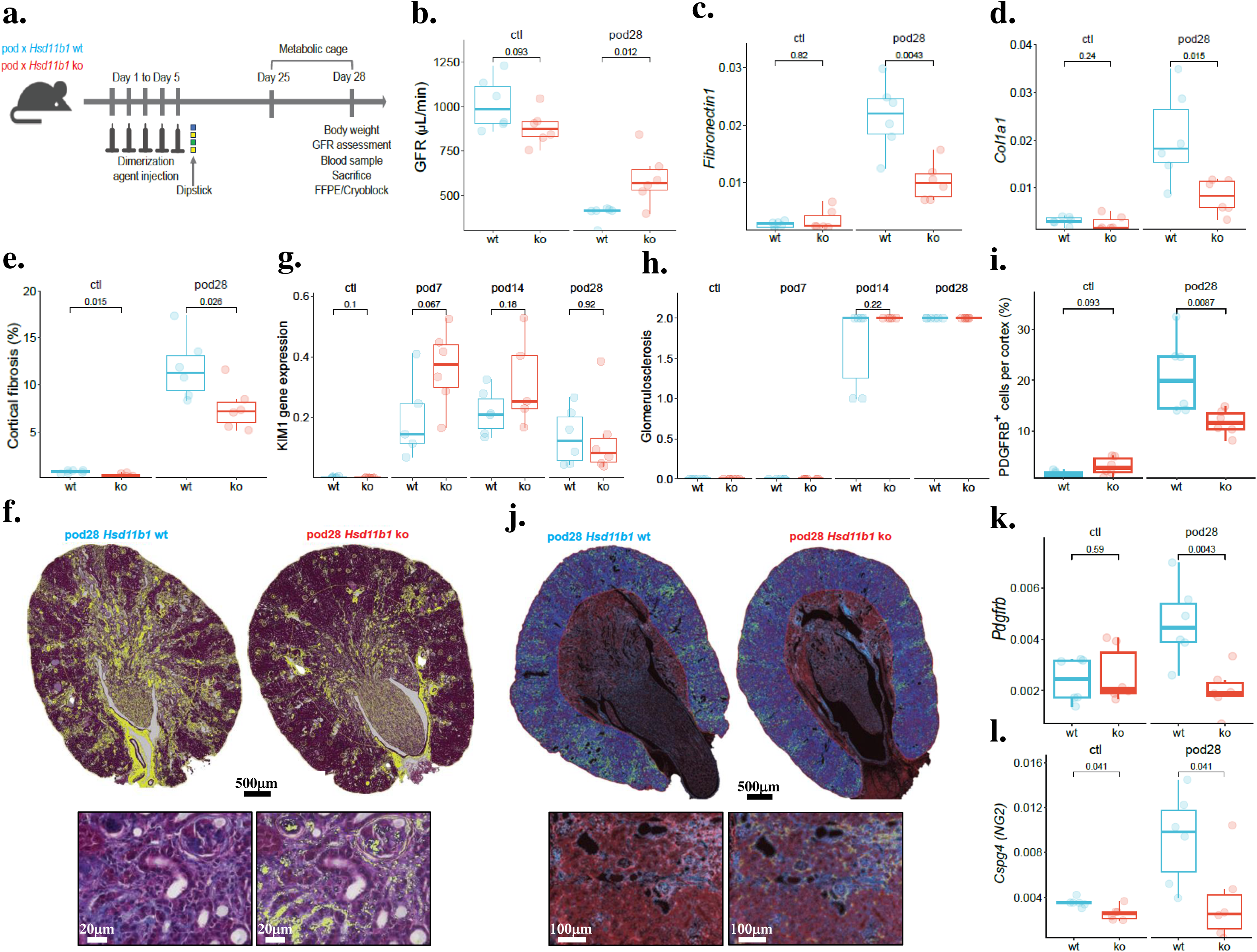
Genetic ablation of *Hsd11b1* ameliorates renal structure and function in CKD. (a) Schematic representation of the experimental strategy (8-12-week male mice, 6 animals per group). FFPE, formalin-fixed paraffin embedded; pod x *Hsd11b1* wt, pod wild-type mice; pod x *Hsd11b1* ko, pod *Hsd11b1* knock-out mice. (**b**) Glomerular filtration rate (GFR) (μL/min) at baseline and after induction of glomerular injury. Ctl, control (uninjured) mice; pod28, POD-ATTAC mice 28 days post-injury; Control (wt), pod *Hsd11b1+/+* mice (wt) (blue); *Hsd11b1 -/-* (ko), pod *Hsd11b1 -/-* (ko) mice (red). (**c and d**) qPCR analysis of gene expression from kidney cortical extracts of wt and *Hsd11b1* ko mice. *Col1a1* (**c**) and *Fibronectin1* (**d**). (**e and f)** Automated quantification of cortical interstitial fibrosis in chronically injured kidney (pod28) from wt and *Hsd11b1* ko animals (**e**) and representative Masson’s trichrome images with overlaid pixel classifier detecting Masson’s trichrome positive tissue area (**f**). (**g**) qPCR analysis of *Havcr1* (KIM-1) gene expression from kidney cortical extracts of wt and *Hsd11b1* ko mice. **(h)** Histological quantification of glomerular sclerosis (glomerulosclerosis). Ctl, control (uninjured) mice; pod7, pod 7 days post-injury; pod14, pod 14 days post-injury; pod28, pod 28 days post-injury. (**i and j**) Representative images with quantification (as % of total cells) (**i**) and object classifier overlay (**j**) of interstitial PDGFRβ+ cells in non-injured (Ctl) and chronically injured (pod28) *Hsd11b1* wt and ko cortices. PDGFRβ, red ; DAPI, blue. (**k and l)** qPCR analysis of gene expression from wt and *Hsd11b1* ko mice kidney cortical extracts. *Pdgfrb* (**k**) and *Cspg4* (**l**).

Correspondingly, mRNA levels of fibroblast markers, including *Pdgfr*β and *Cspg4*^9,16^, were lower in injured kidney cortex of *Hsd11b1* ko mice (Fig.2k-l, Extended data Fig.2o-p). These findings suggest that 11β-HSD1 genetic ablation mitigates TIF in CKD by limiting the expansion of PDGFRβ−positive fibroblasts, regardless of the injury etiology.

### Pharmacological inhibition of 11β-HSD1 mitigates CKD progression

Given the beneficial effects observed with *Hsd11b1* gene deletion on CKD progression, we investigated whether pharmacological 11β-HSD1 inhibition would yield similar results. To address this, we treated pod mice with the potent and selective 11β-HSD1 inhibitor ABT384^18^ , which has previously been tested in clinical trials for diabetes and metabolic syndrome (Fig.3a). First, we confirmed effective *in vivo* inhibition of 11β-HSD1 by observing a significant decrease in the ursodeoxycholyltaurine to 7-oxolithocholyltaurine ratio, a biomarker of pharmacological 11β-HSD1 inhibition^19,20^ (Fig.3b-c). Consistent with our findings in *Hsd11b1* ko animals, pharmacological inhibition of 11β-HSD1 improved GFR, morphometric parameters and reduced TIF estimated four weeks post kidney injury (Fig.3d-g, Extended data Fig.3a). ABT384 also limited fibroblast expansion after injury, as demonstrated by decreased PDGFR-β-positive fibroblast abundance in quantitative immunostaining (Fig.3h-i). These findings indicate that pharmacological inhibition of 11β-HSD1 effectively recapitulates the protective effects of gene deletion on CKD progression.

**Figure 3.**
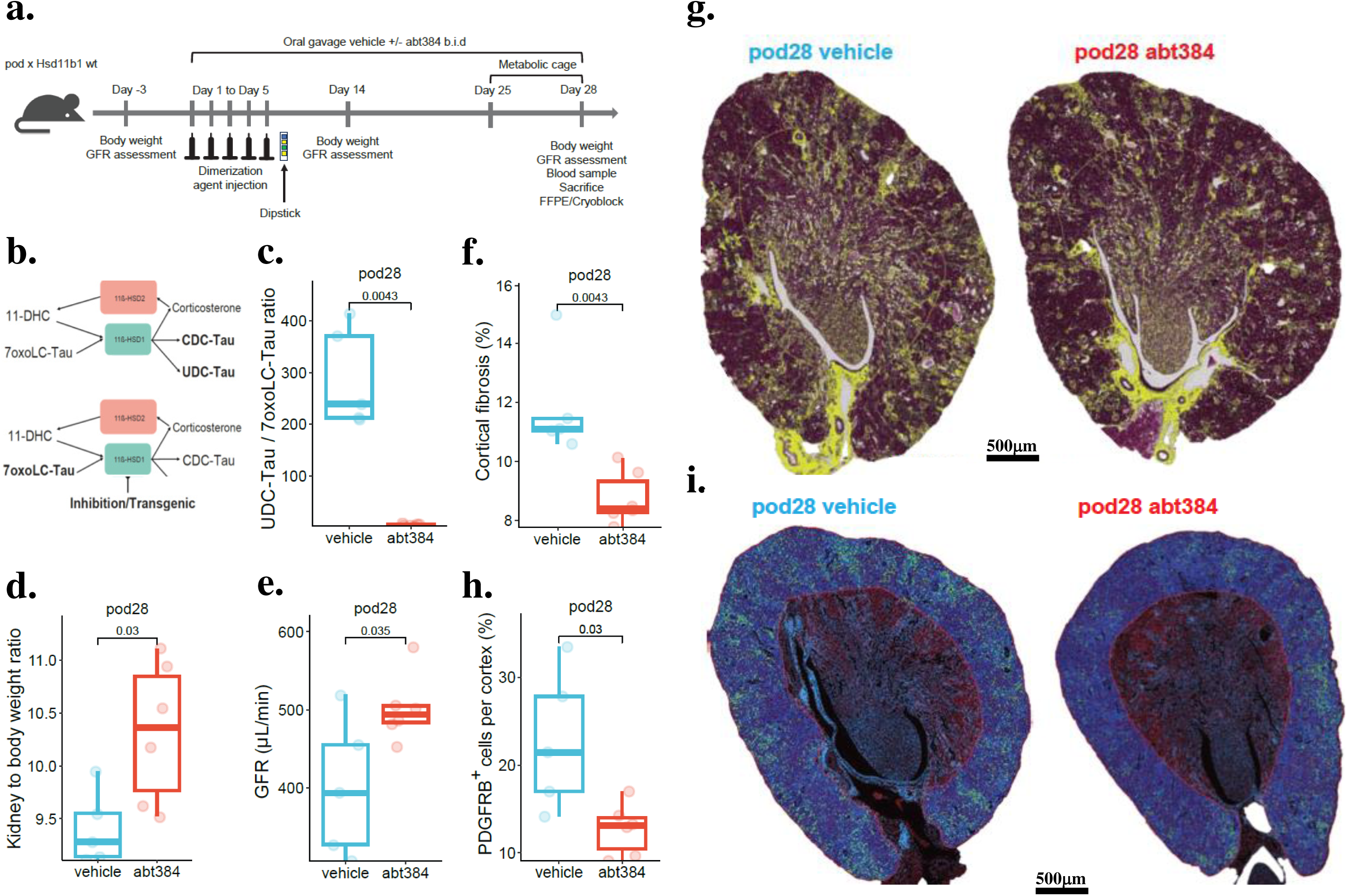
Pharmacological 11β-HSD1 inhibition ameliorates CKD in pod mice. (a) Schematic representation of the *in vivo* pharmacological 11β-HSD1 inhibition experiment. pod x *Hsd11b1* wt, pod wild-type mice (8-12 week old male mice, 5 animals in pod x *Hsd11b1* wt vehicle group, 6 animals in pod x *Hsd11b1* wt ABT384 group). FFPE, formalin fixed paraffin embedded; GFR, glomerular filtration rate. (**b**) Schematic illustration of the effect of 11β-HSD1 inhibition on the ratio of the alternative product ursodeoxycholyltaurine (UDC-Tau) to the substrate 7-oxolithocholyltaurine (7oxoLC-Tau) in plasma. 11-DHC, 11-Dehydrocorticosterone. (**c**) UDC-Tau/7oxo-LC-Tau ratio for vehicle-treated (blue) and ABT384-treated (red) experimental groups. (**d**) Kidney to body weight ratios. (**e**) GFR (μL/min) measured 28 days after glomerular injury. (**f and g**) Automated quantification of cortical interstitial fibrosis in chronically injured kidney (pod28) from pod28 vehicle and ABT384-treated groups (**f**) and representative Masson’s trichrome images with overlaid pixel classifier detecting Masson’s trichrome positive tissue area (**g**). (**h and i**) Representative images (**h**) and quantification (as % of total cells) (**i**) of interstitial PDGFRβ+ cells in pod28 vehicle and ABT384-treated mice. PDGFRβ, red ; DAPI, blue.

### 11β-HSD1 expression is linked to mesenchymal differentiation

In light of the quasi-restricted and persistent expression of *Hsd11b1* in fibroblasts, its significant effect on mesenchymal expansion, and its involvement in TIF progression, we hypothesized that 11β-HSD1 impacts the fibroblast mosaic in CKD.

To explore this, we applied high resolution clustering in our snRNAseq dataset, identifying 4 fibroblastic subpopulations, termed Fib1 to Fib4 (Fig.4a and Extended Data Fig.3b). To annotate these subpopulations, we used scType with a reference atlas of human and mouse fibroblasts using publicly available snRNAseq and scRNAseq datasets generated in this study (Extended Data Fig.3c). Fib1 displayed pericyte characteristics marked by *Myh11* or *Rgs5* while the remaining clusters exhibited fibroblastic and myofibroblastic gene expression profiles. Notably, Fib4 expresses *Gam2* and *Fzb*, markers of late differentiating myofibroblasts^6,21^ (Fig.4b) resembling a fibroblast subtype implicated in tissue fibrosis^4,22^. Based on these gene expression profiles of lineage and functional markers, Fib1 was annotated as pericytes, Fib2 as fibroblasts (Fib), Fib3 as myofibroblasts (MF) and Fib4 as regulatory myofibroblasts (Reg-MF).

**Figure 4.**
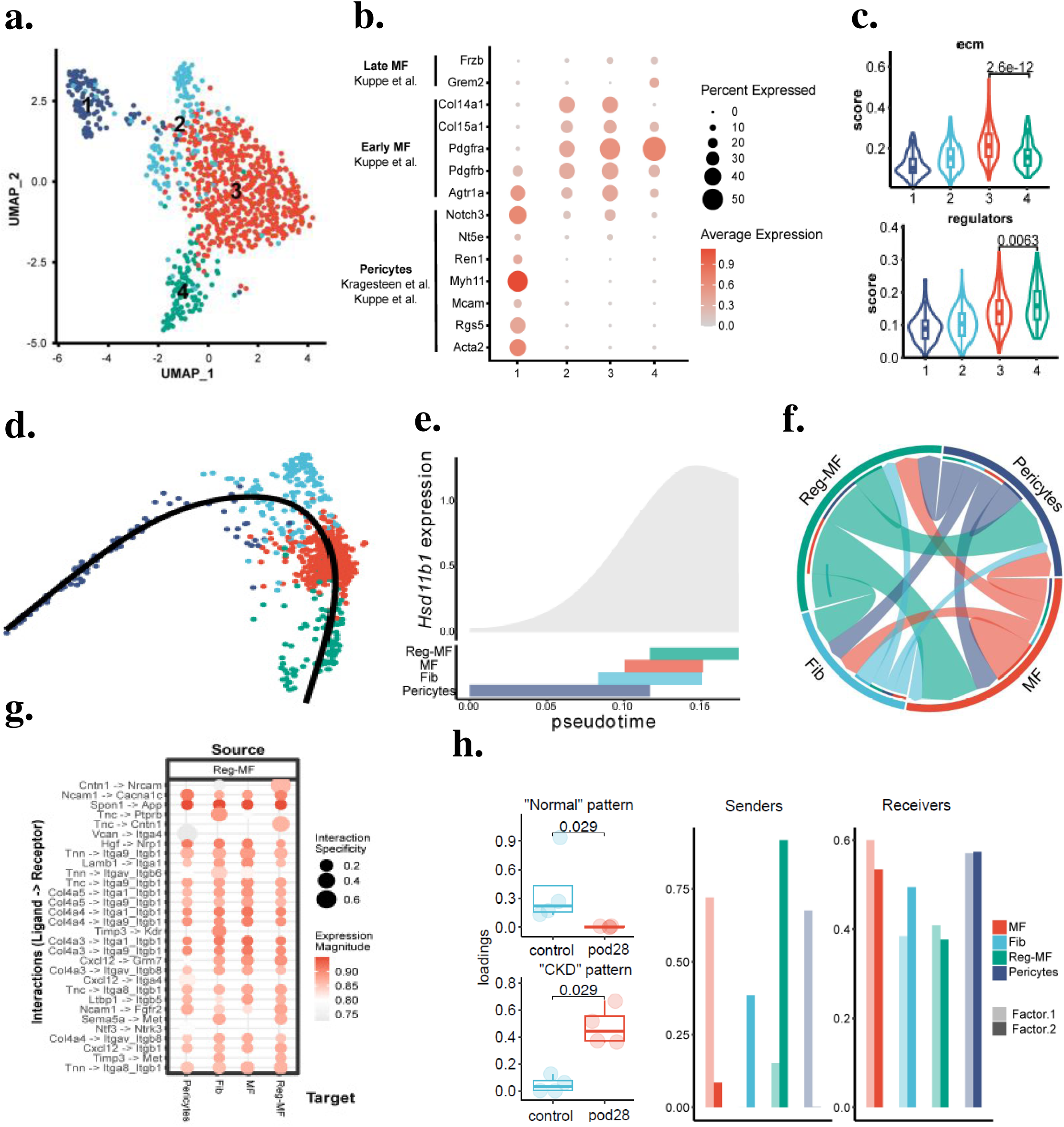
11β-HSD1 participates to myofibroblast heterogeneity. (**a**) UMAP projection and unsupervised clustering of the mesenchymal compartment identifying 4 distinct cell clusters (n = 11 mice). Cluster 1, dark blue; cluster 2, light blue; cluster 3, red; cluster 4, green. (**b**) Dot plot showing the expression of known mesenchymal subpopulation marker genes for each of the 4 clusters identified. Early myofibroblast (MF), late myofibroblast. *Frzb*, frizzled related protein; *Grem2*, gremlin 2; *Col14a1*, collagen alpha-1(XIV); *Col15a1*, collagen alpha-1(XV); *Pdgfra*, platelet-derived growth factor receptor alpha; *Pdgfrb*, Platelet-derived growth factor receptor beta; *Agtr1a*, angiotensin II receptor type 1; *Nt5e*, ecto-5′-nucleotidase (CD73); *Ren1*, renin 1; *Myh11*, myosin heavy chain 11; *Mcam*, melanoma cell adhesion molecule; *Rgs5*, regulator of G protein signaling 5; *Acta2*, smooth muscle alpha-2 actin. (**c**) ECM (left panel) and regulators (right panel) score levels for each of the 4 mesenchymal clusters. (**d**) Pseudotime trajectory embedding of mesenchymal differentiation process for pericytes (originally cluster 1, dark blue), Fib (originally cluster 2, light blue), MF (originally cluster 3, red) and Reg-MF (originally cluster 4, green). (**e**) 11β-HSD1 scaled expression during mesenchymal differentiation. (**f**) Chord diagram showing cell-cell interactions in the mesenchymal compartment. (**g**) Dot plot displaying the magnitude of the expression of the ligands in Reg-MF and the receptors in all mesenchymal clusters. *Cntn1*, contactin 1; *Nrcam*, neuronal cell adhesion molecule; *Ncam1*, neural cell adhesion molecule 1; *Cacna1c*, calcium voltage-gated channel subunit alpha1 C; *Spon1*, spondin 1; *App*, amyloid beta precursor protein; *Tnc*, tenascin C; *Ptprb*, protein tyrosine phosphatase receptor type B; *Vcan*, versican; *Itga4*, integrin subunit alpha 4; *Hgf*, hepatocyte growth factor; *Nrp1*, neuropilin 1; *Itga9_Itgb1*, integrin subunit alpha 9_ integrin subunit beta 1; *Lamb1*, laminin subunit beta 1; *Itga1*, integrin subunit alpha 1; *Itgav_Itgb6*, integrin subunit alpha V_integrin subunit beta 6; *Col4a5*, collagen alpha-5(IV); *Itga1_Itgb1*, integrin subunit alpha I_Integrin subunit beta 1; *Col4a4*, collagen alpha-4 (IV); *Timp3*, tissue inhibitor of metalloproteinase 3; *Kdr*, kinase insert domain receptor; *Cxcl12*, CXC motif chemokine ligand 12; *Grm7*, gremlin 7; *Itgav_Itgb8*, integrin subunit alpha V_integrin subunit beta 8; *Itga8_Itgb1*, integrin subunit alpha 8_integrin subunit beta 1; *Ltbp1*, latent transforming growth factor beta binding protein 1; *Itgb5*, integrin subunit beta 5; *Fgfr2*, fibroblast growth factor receptor 2; *Sema5a*, semaphorin 5A; *Met*, proto-oncogene receptor tyrosine kinase; *Ntf3*, neurotrophin 3; *Ntrk3*, neurotrophic receptor tyrosine kinase 3. (**h**) Identification of specific cell-cell communication programs in control and CKD mice. Left panel shows loadings for each identified program according to CKD status. Middle and right panel show the sender and receiver communication patterns according to their cell type and their factor.

To further characterize the fibroblastic clusters, we interrogated the Matrisome project, a specialized database listing the ensemble of ECM and ECM-associated proteins^23^. According to their mesenchymal nature, all fibroblast clusters displayed high expression of these genes (Extended Data Fig.3d) but with some specificity within the fibroblastic compartment. While Fib3 displayed the highest ECM Core Matrisome activity, Fib4 exhibited the highest score of ECM regulators (Fig.4c), suggesting distinct roles in fibrosis progression. Moreover, each cluster displayed a specific pattern of enriched Gene Ontology pathway including mesenchyme migration (Fib1), collagen binding and fibril aggregation (Fib3, Fib2), and TGFBR activity (Fib4) (Extended Data Fig.3e). Pseudotime analysis supported a differentiation trajectory from pericytes to Reg-MF, with a sequential activation of fibrosis-related genes^24,25^(Fig.4d, Extended Data Fig.3f and g). This trajectory was further strengthened using bulk RNA-seq data from ischemia-reperfusion injury models^26^, where deconvolution showed MF appearing prior to Reg-MF, supporting their place at a later differentiation stage (Extended Data Fig.3h).

To link fibroblast trans-differentiation with our hypothesis that 11β-HSD11 expression in fibroblasts promotes CKD fibrosis, we analyzed *Hsd11b1* expression along the transdifferentiation trajectory. *Hsd11b1* expression increased steadily during this process, peaking in the Reg-MF population (Fig.4e). This aligns with a previous study identifying *Hsd11b1* as one of the mostly expressed genes in *Pdgfr*β positive highly pathogenic ECM-producing fibroblast clusters^6^.

Together, these findings highlight the critical role of 11β-HSD1 in the pericyte-to-myofibroblast transdifferentiation pathway^27^ and suggest that 11β-HSD1-expressing mesenchymal cells are central to the renal fibrotic process in CKD.

### 11β-HSD1 promotes myofibroblast subpopulation diversity and the emergence of a pathogenic subtype in CKD

Cell-cell communication (CCC) analysis using LIANA^28^ identified Reg-MF cells as the primary signaling source, primarily targeting MF and pericyte clusters. (Fig.4f). Detailed ligand-receptor analysis indicated that Reg-MF cells transmit signals via tenascin matricellular proteins, key mediators of tissue remodeling^29^ (Fig.4g). Using Tensor2Cell, we identified context-specific CCC programs, revealing two latent patterns - “normal” and “CKD”. The pattern “CKD” was notably upregulated in Reg-MF cells, supporting their heightened role in signaling during CKD (Fig.4h).

To investigate the spatial relationship between Reg-MFs and other fibroblast subpopulations *in situ*, we performed spatial transcriptomics (Visium). Using the label transfer tool^30^, we deconvoluted spot composition with our snRNAseq dataset as a reference, revealing expected distribution of major cell types across anatomical structures under all experimental conditions (Fig.5a). Consistent with morphometric analysis, fibroblasts primarily localized to injured regions in both wt and ko animals, with reduced abundance in all kidney layers (cortex, outer strip and inner strip of outer medulla) in ko animals (Fig.5b-c, Extended data Fig.4a). Notably, a neighbourhood analysis using *Adamtsl1*, one of the most distinguishing genes of Reg-MF cells (Extended Data Fig.5b), displayed a distinct spatial pattern. Reg-MF tended to aggregate at the “fibroblast border” rather than at the “fibroblast center” (Fig. 5d). This spatial arrangement suggests that Reg-MF cells are strategically positioned at the interface between healthy and fibrotic tissue, where they may regulate fibrosis progression^31^.

**Figure 5.**
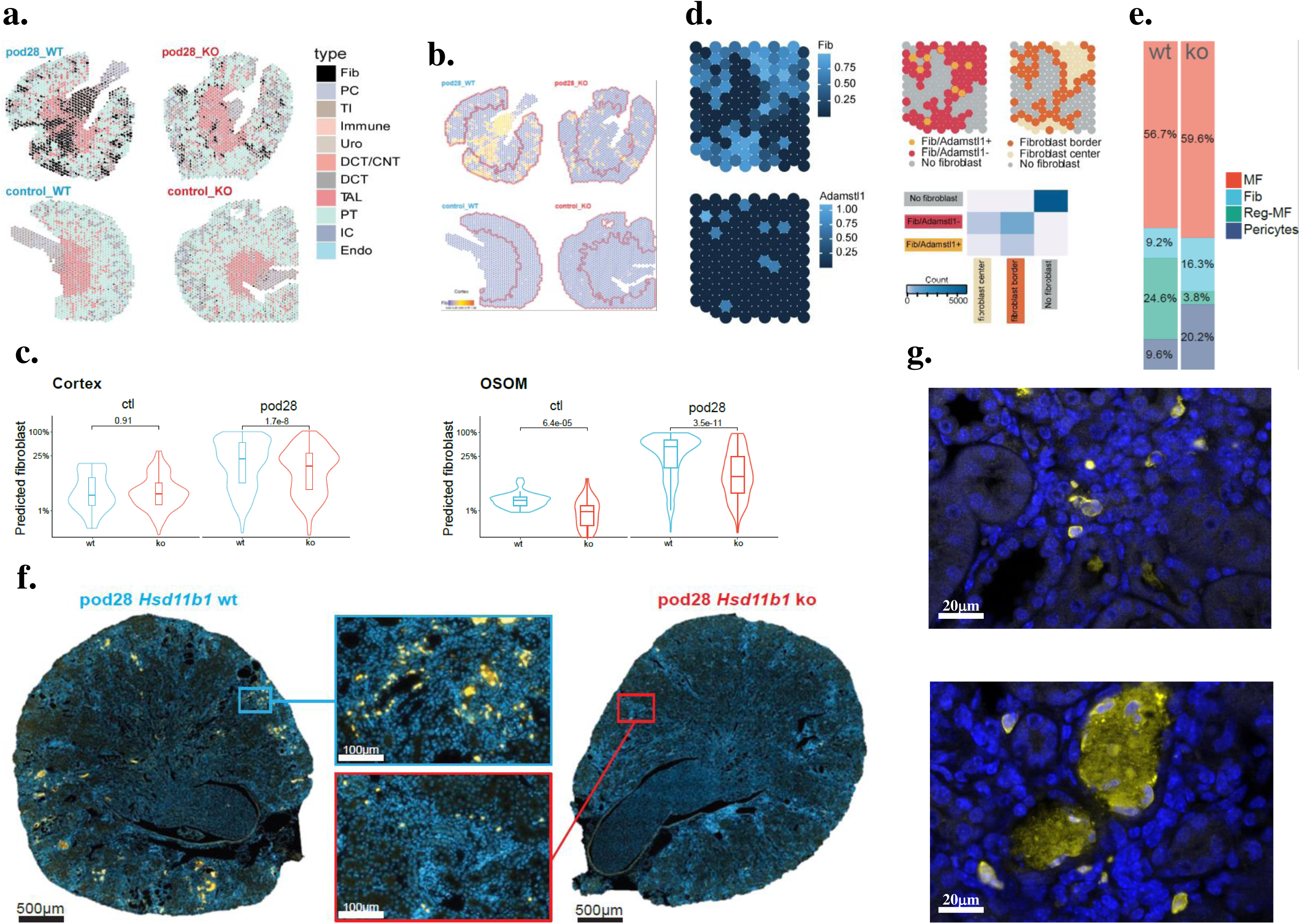
11β-HSD1 promotes the emergence of a myofibroblast pathogenic subtype in CKD. (**a**) Prediction of cell types based on special transcriptomic analyses in kidneys of ctl and pod28 mice *Hsd11b1* +/+ (wt) or *Hsd11b1* -/- (ko). Pie charts show the proportions of transfer scores predicted for cell type annotations using Visium. (**b**) Visium prediction of mesenchymal cluster signatures in the kidneys of control and pod28 mice among genotypes. Cortex boundary is outlined by a red line. (**c**) Violin plot showing the predicted probability of fibroblast detection in each cortical (left panel) and outer strip of outer medulla (OSOM) (right panel) Visium spot from control and pod28 animals. (**d**) Spatial distribution of fibroblast proportion and *Adamstl1* expression per spot for a high-magnification region on Visium (left panel). Spatial distribution of spots with high (gold) or low (red) *Adamtsl1* gene expression relative to fibroblast proportion and differentiation between fibroblast spots based on the boundary between spots with or without fibroblasts (upper right panel). Heat map showing the count of *Adamtsl1* spots with respect to the spatial distribution of fibroblast spots (bottom right panel). (**e**) SnRNAseq relative proportion of each mesenchymal cluster in chronic injured kidney from 2 wt and 2 ko mice. (**f**) Representative immunofluorescence images of ADAMTSL1 in pod *Hsd11b1* wt or ko mice taken 28 days after injury (pod28) (n = 3 per group). ADAMSTL1, gold; DAPI, blue. (**g**) Representative high magnification confocal images of 11β-HSD1/DAPI in kidney cortex injured foci from pod *Hsd11b1* wt mice (n=2). ADAMSTL1, gold; DAPI, blue.

These findings indicate that 11β-HSD1-driven mesenchymal differentiation promotes the emergence of distinct fibroblast subpopulations with specialized roles in fibrosis progression, particularly the Reg-MF cluster, which modulates cellular interactions and tissue remodeling in the fibrotic niche.

### 11β-HSD1 ablation suppresses profibrotic myofibroblast differentiation

To assess whether *Hsd11b1* expression actively drives rather than merely correlates with mesenchymal cell transdifferentiation, we compared fibroblast subpopulations in *Hsd11b*1 ko and wt animals during CKD. Four weeks after pod injury, *Hsd11b1* ko mice exhibited a near-complete absence of Reg-MFs, with a concurrent increase in precursor pericytes and Fib within kidney tissue. This accumulation of myofibroblast precursor cells suggests a differentiation blockade, preventing the transition to Reg-MFs (Fig.5e). Immunofluorescence and confocal analysis confirmed these results, showing strong induction of interstitial ADAMTSL1 labelling in fibrotic areas as well as some intratubular casts from wt animals (Fig.5f-g, Extended data Fig.4c).

A decreased cell cycle proliferation rate is linked to mesenchymal transdifferentiation and is found in several fibrotic disease models^6,32^. We thus assessed whether 11β-HSD1 could impact fibroblast differentiation through an effect on the cell cycle. Using Tricycle^33^, we observed a decrease in the proliferation rate in wt animals along the pseudotime which reaches a minimum during the transition from pericytes to Fib and MF. *Hsd11b1* ko animals maintain a stable proliferation rate (Extended data Fig.4d). Since the cell cycle is tightly regulated by multiple factors, we assessed the effect of 11β-HSD1 on transcription factor expression using the collecTRI-derived regulons dataset^34^. Inhibitor of DNA-binding 2 (ID2) was the sole transcription factor predicted to be significantly altered in *Hsd11b1* ko mice (Extended data Fig.4e). The genetic ablation of *Hsd11b1* fully prevented the strong and early decrease in ID2-dependent target gene expression observed in pod injured wt kidneys (Extended data Fig.4f), suggesting an interaction between 11β-HSD1 and ID2 activity during CKD progression.

We next investigated whether pharmacological inhibition of 11β-HSD1 recapitulated these effects. Deconvolution analysis of bulk RNAseq data from ABT384-treated and untreated pod kidneys revealed a selective reduction in Reg-MF abundance, while other fibroblast subpopulations remained unaffected (Extended data Fig.4g, Extended data Fig.4h). As expected, there was a significant association between GFR and abundance of myofibroblast population (Extended data Fig.4i). These findings suggest that 11β-HSD1 inhibition exerts its protective effect primarily by curbing Reg-MF differentiation rather than MF proliferation.

### Reg-MF cluster is conserved in the Human kidney and predicts CKD progression

To evaluate the translational relevance of our findings, we assessed the presence of the Reg-MF subpopulation in kidneys of CKD patients. Leveraging the largest available snRNAseq dataset of human kidney tissue, which encompasses 23 patients (10 with CKD and 13 control)^35^, we mapped mouse fibroblast cluster signatures onto the human dataset. Through integration of these data into a newly computed UMAP, we identified pericytes, Fib, MFs, and Reg-MFs as distinct subpopulations within the fibroblast compartment, with minimal overlap between them (Fig.6a). This finding supports the conservation of these fibroblast subpopulations across species and suggests that Reg-MFs may play similar functional roles in human CKD.

**Figure 6.**
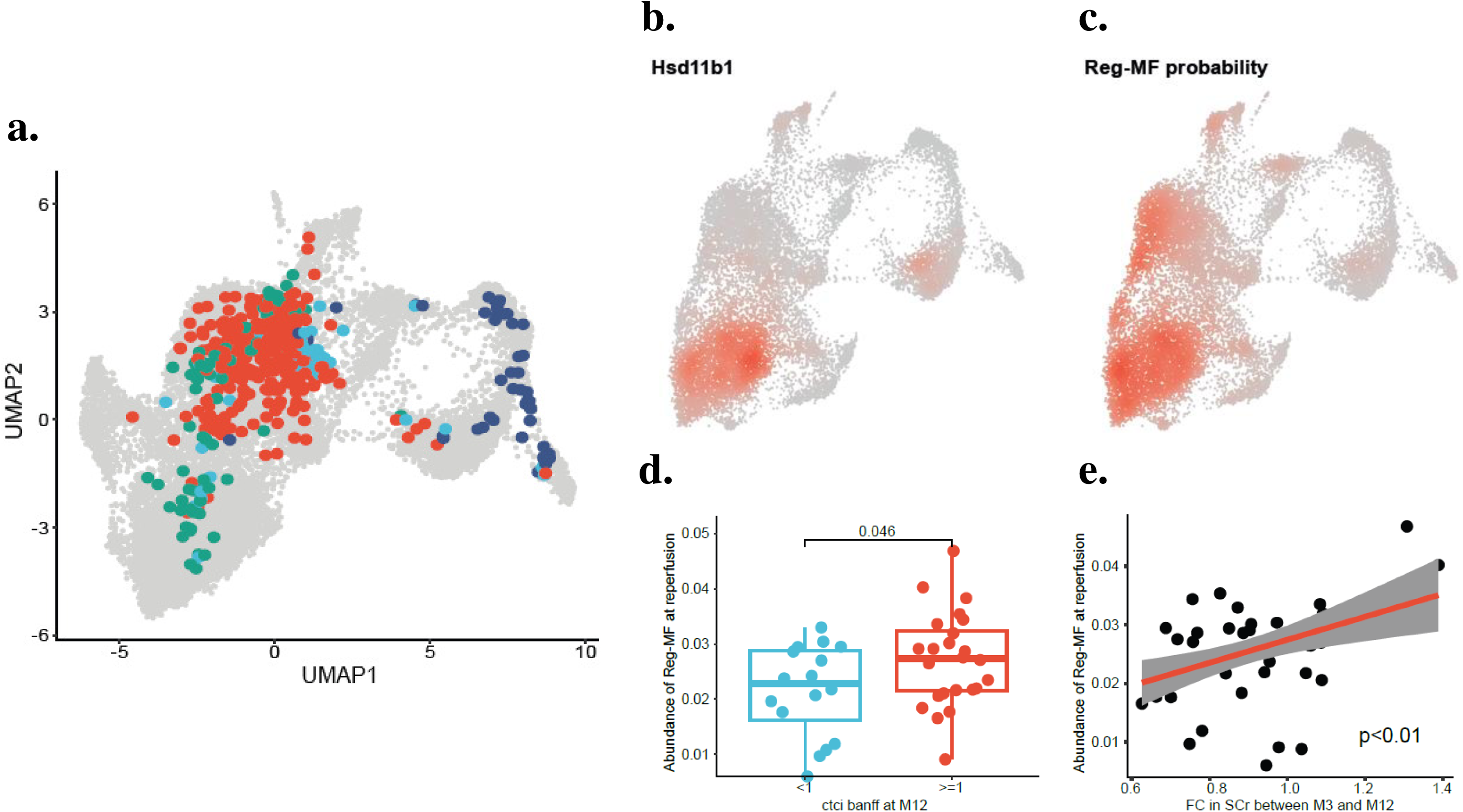
Reg-MF cluster predicts CKD progression in Human. (**a**) UMAP projection of the four mesenchymal subpopulations onto human kidney snRNAseq datasets. Fib, fibroblast; MF, myofibroblast; Reg-MF, regulator-myofibroblast. (**b and c**) Predicted *HSD11b1* expression (**b**) and Reg-MF signature (**c**) onto human kidney snRNAseq datasets. (**d**) Comparison of the estimated abundance of the Reg-MF cluster in transplanted human kidneys combined tubular atrophy (ct) and interstitial fibrosis (ci) pathological Banff Classification at reperfusion and one-year after kidney transplantation (n=42). (**e**) Correlation between estimated abundance of the Reg-MF cluster in transplanted human kidneys at reperfusion and serum creatinine change between 3 and 12 months (n=42).

Consistent with our observations in mice, we noted similar patterns of *HSD11B1* expression in renal fibroblastic populations and a similar distribution of Reg-MF based on transfer learning predictions (Fig.6b-c). To explore whether Reg-MF abundance correlates with clinical outcomes, we analyzed our cohort of renal bulk RNAseq data from 42 kidney allograft recipients at the time of transplantation reperfusion^36^. Cell-type deconvolution revealed that higher Reg-MF abundance at reperfusion was positively associated with more severe chronic histological lesions one year post-transplant (Fig.6d). Furthermore, Reg-MF abundance at reperfusion was negatively correlated with serum creatinine at 3 months post-transplant (Fig.6e), indicating the potential of Reg-MF abundance as a prognostic marker for CKD progression. Interestingly, no perioperative clinical variable correlated with Reg-MF abundance at reperfusion (Supp Table 1).

Together, these data underscore the cross-species conserved role of Reg-MF in promoting CKD progression and highlight the translational relevance of our findings.

## Discussion

CKD represents a significant global health burden, with limited therapeutic options beyond targeting the renin– angiotensin–aldosterone system. TIF is a hallmark of CKD progression, yet direct antifibrotic therapies remain elusive^3,37^, largely due to an incomplete understanding of the kidney fibrogenic microenvironment and the heterogeneity of myofibroblast populations^6,38^. Herein, we demonstrated that genetic deletion or pharmacological inhibition of 11β-HSD1 markedly attenuates interstitial fibrosis and improves renal function in CKD models. These protective effects are achieved by limiting the expansion of the Reg-MF population, a highly pathogenic subset of myofibroblasts. Our study places 11ß-HSD1 in the driver seat of TIF progression and trans-differentiation from pericyte progenitors to myofibroblast, and demonstrates that inhibition of the intracellular activation of glucocorticoids, which limits fibrogenesis, is a potential treatment option in CKD.

Since 11ß-HSD1 regulates the local levels of active glucocorticoids, our findings support the idea that kidney fibrosis is stimulated by endogenous glucocorticoids. On one hand, this observation may appear surprising since treatment with exogenous GC have been used clinically for the treatment of fibrotic diseases in other indications, particularly for idiopathic lung or cystic fibrosis^39^, although their clinical efficacy remains to be clearly demonstrated^40^. The anti-inflammatory action of pharmacological glucocorticoids like prednisone/prednisolone and dexamethasone through glucocorticoid receptor (GR)-mediated inhibition of pro-inflammatory mediators released by immune cells was the main intended mechanism of action proposed to rationalize the potential benefit of glucocorticoids treatment in fibrotic diseases, but the actual effect of glucocorticoids concentration on fibroblast populations transactivation/transrepression remained underexplored. On the other hand, more recent observations showed that glucocorticoids administration can enhance the pro-fibrotic effect of transforming growth factor (TGF)-β on fibroblasts which is fully consistent with the profibrotic activity of 11ß-HSD1 observed in our study^41,42^. Although downregulation of the corticosterone response observed in fibroblasts of *Hsd11b1* ko animals strongly supports a pro-fibrotic activity of 11β-HSD1-mediated endogenous GC through GR activation in fibroblasts, the involvement of the MR cannot be excluded since endogenous GC exposure can also result in MR activation^10^. In line with this idea, the MR is described as a promoter of fibrotic remodeling in heart diseases and CKD^43,44^. Recent therapeutic assays indicate that MR antagonists displayed some efficacy in retarding disease progression in CKD patients^45^ but enthusiasm is dampened by potentially life-threatening side effects dominated by hyperkaliemia. Further investigations are warranted to uncover whether the protective role of 11β-HSD1 inhibition is mediated through GR or/and MR. Nevertheless, the role of 11β-HSD1 in myofibroblast trans-differentiation is consistent with a cell proliferation blockade often observed in response to GR activation^46^ and seems to be restricted to organs with pericyte only-derived myofibroblast transdifferentiation, as suggested by conflicting results reported for 11β-HSD1 deletion in the liver with at least two myofibroblast precursor source, i.e. pericyte-like stellate cells and liver progenitor cells ^47,48^.

We showed that genetic *Hsd11b1* deletion or pharmacological inhibition of 11β-HSD1 result in the quasi-absence of a tissue fibrosis-promoting regulatory myofibroblast subpopulation and dramatically improve morphology and function in CKD models. This myofibroblastic subpopulation was also found in human kidneys and its abundance was predictive of CKD progression. In addition, these myofibroblasts were characterized by the specific expression of ADAMTSL1, an ADAMTS-like protein without catalytic activity, viewed as an inhibitor of ADAMTS metalloproteinases which participate in ECM degradation^49^. Consistently, Adamts2 ko mice displayed increased interstitial fibrosis after aortic banding^50^. Interestingly, ADAMTSL1 expression has been linked to maladaptive tissue repair and cardiac fibrosis in human hearts^51^. Taken together with our findings on kidney fibrosis, current knowledge suggests that ADAMTSL1 could play a direct role in maladaptive tissue repair. In addition, detection of highly pathogenic regulatory myofibroblasts through ADAMTSL1 expression may help to select patients with bad prognosis who may best benefit from 11β-HSD1 inhibition.

In conclusion, this study positions 11β-HSD1 as a pivotal regulator of the fibrogenic niche in CKD, driving the differentiation of a pathogenic regulatory myofibroblast subpopulation. Targeting this enzyme may offer a novel and clinically relevant strategy to combat CKD progression. Given the availability of 11β-HSD1 inhibitors developed for other indications^52,53^, there is a clear and timely opportunity to translate these findings into impactful therapies for CKD patients.

## Methods

### Animal experiments

The *Hsd11b1* ko mouse model was generated by the Transgenic Core Facility of the University of Geneva using CRISPR/Cas9 technology. Briefly, after *in vitro* fertilization, oocytes from B6D2F1 mice were injected with Cas9 protein and guide RNA (gRNA) targeting *Hsd11b1*. Two gRNAs were injected to induce the excision of a 552 bp sequence including exons 4 and 5 and generating a frameshift (gRNA 1 and gRNA 2; Supplementary Table 2). To validate the excision, mice were genotyped using PCR primers targeting either side of exons 4 and 5 (Fw1 and Rv1 primers; Supplementary Table 2). The PCR product was 1080 bp long for the wt allele and 552 bp for the ko allele. B6D2F *Hsd11b1* ko mice were then backcrossed to C57BL/6 for 10 generations. The glomerular CKD mouse model pod was previously described^54^. Genotyping of transgenic mice was performed by PCR analysis (Fw1 and Rv1 primers; Supplementary Table 2). Briefly, apoptosis of glomerular podocytes was induced by five times intraperitoneal injections of 0.2 μg/g body weight of a chemical dimerizer (AP20187, Takara Bio Inc. Kusatsu, Japan) in 8-12 weeks old male C57BL/6J pod/*Hsd11b1* +/+ and C57BL/6J pod/*Hsd11b1* -/- transgenic mice. Proteinuria was confirmed by a urine dipstick test (Combur test HC, Roche, Switzerland) at day 5. Mice were sacrificed at day 28 after the first injection. Unilateral ureteral obstruction (UUO) was performed as previously described^55^. Briefly, 8-12 week-old male mice (C57BL/6J *Hsd11b1* +/+ and C57BL/6J *Hsd11b1* -/-) were anesthetized with isoflurane and underwent left unilateral ligature of the ureter. Mice were sacrificed at day 7 after UUO. For pharmacological 11β-HSD1 inhibition, treatment of mice was initiated in the morning preceding the first dimerizer injection until the night preceding the euthanasia. 11β-HSD1 inhibitor ABT384 or vehicle (0.5% methylcellulose, 0.2% Tween-80) were administered orally twice a day (7 a.m. and 7 p.m.) at 10 mg/kg in 200 μL (with a PTFE feeding needle, 20 G, diameter 1.5 inches, length 1.9 mm).

### Transcutaneous GFR measurement

The glomerular filtration rate (GFR) was measured by FITC-sinistrin fluorescence decay as previously described^56^. FITC-sinistrin was infused into the tail vein and fluorescence recording was analyzed with MPD Lab software (Mannheim Pharma and Diagnostics, Germany). GFR was calculated using the appropriate formula^57^ and expressed as μL per minute. For quantitative PCR (qPCR) total RNA was extracted from the renal cortex using a NucleoSpin RNA II RNA extraction kit following the manufacturer’s instructions (Macherey-Nagel, Germany). mRNA was reverse transcribed using qScript cDNA Super Mix (Quanta Biosciences, USA). Ten nanograms of cDNA was used for the qPCR reaction with SYBR Green Master Mix (Applied Biosystems, USA), and the reaction was performed in duplicates using the QuantStudio 5 and StepOne Real-Time PCR System (Thermo Fisher Scientific, USA). The data were normalized to the reference gene, ribosomal protein lateral stalk subunit P0 (Rplp0), and results are expressed as 2^-DDCT.

### Blood pressure measurement

Blood pressure (BP) was measured using the noninvasive CODA system (Kent Scient. Corp., Torrington, CT). Mice were acclimated to the system over 5 days (2:00 PM) and measurements done over 4 consecutive days (2:00 PM). Mice (4 mice simultaneously) were placed in pre-warmed holders on a heating platform in order to maintain the tail temperature between 32° and 35°C. After 5 cycles of acclimation, blood pressure was measured over 20 successive cycles. For each cycle, the occlusion tail cuff is inflated to momentarily impede blood flow. During the slow deflation, a second tail cuff incorporating Volume Pressure Recording (VPR) technology, measures the physiological characteristics of the returning blood flow. Systolic BP is measured automatically at the first appearance of tail swelling and diastolic BP is measured automatically when the rate of swelling stops increasing in the tail. Total duration of the test was 15 minutes.

### Proteinuria measurement

Proteinuria was measured using the Indiko system (Thermofisher Scientific, Waltham, USA). 24h-urine was collected the morning preceding the transfer to the metabolic cage, quantified and stored at -80°C. Protein measurement were obtained by photometric determination using molybdenum and pyrogallol red stains.

### Histology and immunofluorescence imaging

PFA-fixed kidney tissues were embedded in paraffin following 2 h fixation with 4% paraformaldehyde (PFA) in PBS. Sections were cut at a thickness of 5 µm and stained with hematoxylin/eosin or Masson’s trichrome. For immunofluorescence, heat-induced antigen retrieval was applied on PFA-fixed tissues either by microwave or pressure cooker in pH 6 citrate or pH 9 EDTA buffer. Tissue sections were washed in TBST (0.1% Tween20 in TBS), permeabilized with 0.2% Triton X-100 in TBST, blocked with 2% bovine serum albumin in TBST, incubated overnight at 4°C with primary antibodies, detected with species-specific secondary antibodies coupled to Alexa Fluor 488 or 647 (Jackson ImmunoResearch, USA) for 1 h, and incubated with DAPI for 10 min at room temperature. The following antibodies were used: 11β-HSD1 (Sigma, no. HPA047729, 1:50), PDGFRβ (Invitrogen, no.MA5-15143, 1:50), PDGFRβ (Abcam, no.AB69506, 1:50), ADAMTSL1 (Sigma, no.HPA057437, 1:50). Sections were mounted with Vectashield (Vector Laboratories, USA). All images were acquired on a Zeiss Axio Scan Z1 and 3D Histech Panoramic 250 flash.

### Confocal imaging

PFA-fixed kidney tissues were embedded in paraffin following 2 h fixation with 4% PFA in PBS. Sections were cut at a thickness of 5 µm. Slides were prepared from pod *Hsd11b1* wt animals (n=2). Heat-mediated antigen retrieval was performed using a pressure cooker (2x10 min; low pressure) in pH 6 citrate buffer. After washing in TBST, the sections were permeabilized in 0.2% Triton X-100 in TBST and blocked with 2% BSA in TBST. The sections were incubated with the primary anti-ADAMTSL1 antibody (Sigma, ref. HPA057437; 1:50), 11β-HSD1 (Sigma, no. HPA047729, 1:50) and PDGFRβ (Invitrogen, no.MA5-15143, 1:50) overnight at 4°C, detected with a species-specific secondary antibody conjugated to Alexa Fluor 647 for 1 hour and incubated with DAPI for 10 min at room temperature. The slides were mounted using Vectashield and image acquisition was performed using a Leica Stellaris 5 confocal microscope.

### Kidney fibrosis quantification

Digitalized Masson’s trichrome stained sections were further processed and analyzed using QuPath (0.4.0) open-source software^58^. A script based on vectorial color-deconvolution in RBG space was generated as previously described^59^. The cortical area was delineated using glomeruli and arcuate vessels as cortical boundaries and glomerular areas were removed in a blinded manner on each section by an experienced nephropathologist. The extent of interstitial fibrosis was expressed as a percentage of affected cortical areas.

### Immunofluorescent image quantification

Interstitial cells expressing PDGFRβ on mouse immunofluorescent sections were quantified using QuPath (0.4.0) open-source software as previously described^60^. Briefly, the cortex was manually annotated as above, and cells were detected using the DAPI algorithm-based “Detect Cells” tool, using the optical density sum to segment nuclei. Cells were then classified with the “Train Object Classifier” tool using a neuronal network trained on 20-40 annotations of each cell class (interstitial PDGFRβ positive or negative cell) for each experimental condition (control wt and *Hsd11b1* ko animals; pod wt and *Hsd11b1* ko animals). Object classifier algorithm was then applied to each slide of the dedicated condition. Results are expressed as a percentage of interstitial PDGFRβ positive cells per all cortical nuclei.

### 3D spatial analysis

Images were acquired using a Leica Stellaris 5 confocal microscope using 20x and 40x objectives. Raw imaging data were processed using Imaris software for 3D reconstruction and Adobe Illustrator. 3D confocal images of 11β-HSD1 expressing cells were manually annotated using Imaris 10.0.1 (Andor Technology Limited, UK) with spots centered on their nuclei indicated by the DAPI channel. Spatial analysis was conducted with MATLAB R2023a (The MathWorks Inc., USA). Nearest Neighbor analysis was carried out by measuring all pairwise Euclidean distances between each cell, retaining the shortest distance. Ripley’s K analysis performed by using the methodology as previously described^61^. Briefly, Ripley’s K function is defined as:

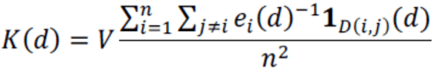

Where V is the considered volume, n is the number of cells in the volume, 𝑒i(𝑑) is the edge correction, 𝟏D(i,j)(𝑑) is the indicator function and finally 𝐷(𝑖, 𝑗) is the Euclidean distance from the cell i to j. Under complete spatial randomness, the expected value of K, is:

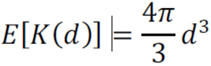

Ripley’s K function subtracted by its expected value. Green and bold red lines indicate the respective average of control and pod28 mice, the filled areas represent the 95% confidence intervals, the horizontal black dashed line indicates a complete spatial randomness.

### RNA-sequencing

For bulk RNAseq, 11 kidneys (n = 5 and n = 6 for vehicle-treated and ABT384-treated mice, respectively) were included. RNA extraction, library preparation and sequencing were performed as previously described^62^. Briefly, the total RNA was extracted from kidney tissues, according to the manufacturer’s instruction. The RNA quantification was performed with a Qubit fluorimeter (Thermo Fisher Scientific, Illkirch, France) and the RNA integrity was assessed with a Bioanalyzer (Agilent Technologies, Les Ulis, France). The TruSeq mRNA stranded kit from Illumina was used for library preparation with 700 ng of total RNA as input. The library molarity and quality were assessed with the Qubit and Tapestation using a DNA high sensitivity chip (Agilent Technologies). The libraries were pooled at 2 nM and loaded for clustering on a Single-read Illumina Flow cell for an average of 25 million reads per sample. Reads of 100 bases were generated using the TruSeq SBS chemistry on an Illumina NovaSeq 6000 (Illumina, Evry, France). The quality control was then performed with FastQC and the reads were mapped to the GRCm39 genome using STAR software. Tables of counts were then generated with HTSeq. Raw counts were further processed as an input for cell-type deconvolution (described below).

### Single-nucleus RNAseq and data analysis

All raw FastQ files were processed using the 10x Cell Ranger counts pipeline^63^ with default settings. This pipeline maps FASTQ reads to a reference genome, filters and counts unique molecular identifiers (UMIs) against a gene reference, estimates the number of cells in the sample, and produces a matrix of gene counts for each cell. Since version 7.0.0 of the pipeline was employed, both intronic and exonic reads were counted. Alignment and counting were performed against the mm10-2020-A mouse reference, as provided by the 10x file server. Estimated cell numbers ranged between 3,148 and 8,401 cells, with an average mean reads-per-cell ratio of 26,193 and an average median genes per-cell ratio of 1,652. One of the two ko pod7 samples deviated, with an estimated 22,985 cells, a mean reads-per-cell ratio of 6,965, and a median genes-per-cell ratio of 817. This outlier was excluded from downstream analysis.

### Cell clustering analysis

All filtered barcode matrices (matrices of gene counts per cell) were loaded into R using the Seurat package version 4.3.0.166 for graph-based cell clustering analysis and visualization. Low-quality cells, putative doublets, and cells undergoing apoptosis were filtered out if they either contained more than 500 or fewer than 7,000 unique detected RNA genes, or had fewer than 1,000 unique molecular identifier (UMI) counts. Before filtering, multiplet identification was conducted using the DoubletFinder algorithm^64^. To correct potential ambient RNA contamination, SoupX algorithm^65^ was also employed accordingly. After correction, datasets were further processed using the SCTransform function to normalize and stabilize gene expression variance across cells, using a Generalized Linear Model with a Gamma-Poisson (glmGamPoi) distribution to address overdispersion. Additionally, the per-cell percentage of mitochondrial transcripts was regressed out during normalization. The SCTransform procedure also included feature selection to identify the 3,000 most variable genes across cells. Following normalization, feature selection, and scaling, Principal Component Analysis (PCA) was performed on each dataset using the RunPCA function. The first 50 principal components (PCs) were used for graph-based clustering using the Louvain algorithm, at a resolution of 0.6 (executed by FindNeighbors and FindClusters functions). For visualization, the UMAP method was applied to the first 50 PCs, offering a two-dimensional representation of single-cell datasets for visual interpretation of cellular clusters and states based on gene expression. Clusters with heterogeneous profiles were further refined by sub-setting and rerunning the afore mentioned analysis pipeline. If these clusters remained heterogeneous after refinement, they were excluded from further analysis. This process ensured the identification of stable cell clusters across all individual samples.

### Data integration

To project all samples into a shared dimensional space, we employed the reciprocal PCA (rPCA) strategy provided by the Seurat R package. Briefly, 3,000 integration features were chosen to identify integration anchors between datasets. Consistent with the single-sample analysis, the SCTransform method was utilized for data normalization. The first 30 components of rPCA were designated for the neighbor search space, using 20 neighbors to select anchors between datasets. To reduce memory usage during pairwise anchoring in the integration process, the first replicate of each genotype and condition was used to form a reference set for integration. The integration process generated a corrected count matrix for the 3,000 anchor features, that was used as input for another round of PCA. The first 50 PCs were then employed for graph-based clustering using the Louvain algorithm. The integrated low dimensional space was then visualized by UMAP, using again the first 50 principal components. Upon visually examining the cellular clusters, non-stable clusters were divided and either retained or excluded.

### Cell cluster annotation

Cell clusters were annotated with respect to their cell-type origin, using previously characterized markers^26,66,67^ and cell types, including: B-cells (*Igha*), connecting tubule - CNT (*Slc8a1*, *Scnn1g*, *Calb1*, *Egfem1*), distal convoluted tubule – DCT (*Slc12a3*, *Slc8a1*), endothelial (*Flt1*, *Pecam1*, *Egfl7*), fibroblasts (*Prkg1*), intercalated cells type A - ICA (*Aqp6*, *Kit*, *Atp6v0d2*, *Atp6v1g3*), intercalated cells type B - ICB (*Atp6v0d2*, *Slc26a4*, *Atp6v1g3*, *Slc4a9*), immune (*Cd247*, *Ptprc*), leukocytes (*Tprc*, *Cd74*, *Wdfy4*), macrophages (*Ptprc*, *Itgam*, *Cd74*), macula densa - MD (*Slc12a1*, *Tmem207*, *Enox1*, *Thsd4*, *Nos1*), medullary thick ascending limb - MTAL (*Slc12a1*), myofibroblasts (*Col1a1*, *Col3a1*), principal cells – PC (*Scnn1b*, *Scnn1g*, *Aqp2*, *Aqp4*), parietal epithelial cells - PEC (*Ncam1*), podocytes - POD (*Nphs1*, *Nphs2*), proximal tubule - PT (*Slc34a1*), proximal tubule segment 1 - PT-S1 (*Lrp2*, *Slc13a3*, *Slc5a12*), intersegment between proximal tubule segment 1 and segment 2 - PT-S1S2 (*Cyp2e1*, *Slco1a1*), proximal tubule segment 3 - PT-S3 (*Lrp2*, *Slc16a9*, *Slc27a7*, *Cyp7b1*), stromal (*Itga8*, *Pdgfrb*, *Hsd11b1*, *Cfh*), thick ascending limb - TAL (*Slc12a1*, *Tmem207*, *Umod*, *Enox1*), thin limb-tL (*Epha7*, *Aqp1*, *Jag1*, *Cdh6*, *Cdh13*), and urothelium - Uro (*Slc12a2*, *Upk1b*, *Psca*). Each sample was annotated separately in a cluster-specific manner. The annotation information was then transferred to the rPCA integrated space and was refined by using the same procedure on the rPCA-derived clusters. This procedure resulted in 59,531 high-quality annotated nuclei. We then used the fibroblast compartment previously identified and rerun RunPCA, RunUMAP, FindNeighbors and FindClusters commands with default parameters. The optimal clustering resolution was selected using a clustering tree and looking for stable clusters. Gene markers for each cluster were found using the FindAllMarkers function. Gene expression levels were displayed on the UMAP using the Nebulosa package based on kernel density estimation. Extracellular Score Matrix and Regulators scores were calculated as previously described^6^ using the core matrisome dataset^68^. Identification of fibroblast subtypes was performed using known gene^6,21^ and the computational method ScType^69^, which enables a fully automated cell-type identification. For this last approach, we built a comprehensive cell marker database including 21 renal single-cell or single-nucleus RNAseq datasets, as background information. Pathway enrichment analysis was performed using the Singscore method^70^, and top pathways across clusters were identified using the irGSEA.integrate.R command from the irGSEA package, which is a wrapper for the Seurat::FindMarkers command. P values were calculated through Wilcoxon’s test and corrected for multiple tests with the Bonferroni method. To study cell-cell communication, we used the Liana algorithm^28^ that infers communications between cells as a consensus among various methods. We applied default parameters to theLiana_wrap command, and aggregated communication scores calculated with the following methods: natmi, connectome, logfc, sca, and cellphonedb. To further decipher cell-cell communication programs across conditions, we combined two tools, LIANA algorithm and Tensor-cell2cell^71^. The analysis was conducted according to the pipeline recently described^72^. We used the magnitude_rank as communication scores. Transcription factor activity was predicted by running the Univariate Linear Model implemented in the decoupleR package. The CollecTRI regulons was used as an input of prior knowledge resource. Pseudotime was inferred on PHATE reduction^73^ using Slingshot^74^ with cluster Fib1 (pericytes) defined as the starting point. Gene expression and transcription factor activity among pseudotime were further fitted using tradeSeq by running the fitGAM function, after having determined the optimal number of knots through the evaluate K function. We took advantage of the “Condition” parameter to fit two different models per gene or transcription factor, corresponding to the genotype status. Genes and transcription factors associated with pseudotime and displaying a different pattern of expression across genotypes were identified by the associationTest command, with a false discovery rate threshold setting at 0.1. Geneset enrichment analysis was performed on the genes significantly associated with pseudotime using the fgseaMultilevel function from the fgsea package. To assess the proliferation rate, we firstly inferred the cell cycle position θ using the tricycle algorithm^33^. We then discretized all cells along into two bins corresponding to actively proliferating (0.25π<θ<1.5π; S/G2/M) or non-proliferating (G1/G0), as previously described^33^. For KPMP cells projection, we downloaded the Seurat object including snRNAseq dataset from 29 patients 10 of whom had CKD and 13 controls, excluding patients with AKI from atlas.kpmp.org (downloaded in 2023.04). We reprocessed all the data from the beginning to ensure homogeneity with our own dataset. Firstly, we split the dataset by patients and performed SCT normalization. We then integrated the data together using the RPCA method from Seurat. Once merged, we subdivided fibroblast and vascular smooth muscle cell types according to the subclass.l1 metadata column. Finally, we ran the RunPCA and RunUMAP commands with default parameters. We then used the FindTransferAnchors and the MapQuery commands using the previous KPMP object as the reference and our own pod mice as the query dataset. To ensure compatibility, we converted gene symbols from the KPMP dataset into their mouse orthologs using the convert_human_to_mouse_symbols command from the nichenetr package. To ensure that cell identified into our dataset were also found in the KPMP dataset, we did not directly project them onto the KPMP UMAP. As suggested^75^, we computed a new UMAP (de novo visualization) after merging the reference and query datasets.

### Cell deconvolution

We performed cell-type deconvolution of bulk-RNAseq datasets using MuSiC algorithm^76^. As the reference snRNAseq dataset, we used the full Seurat object including all renal cell types from wild-type animals. Deconvolution was applied firstly on two mice bulk-RNAseq datasets. The first one included mice undergoing acute kidney injury following ischemia reperfusion injury and sacrificed at various time points up to 12 months^77^. This dataset was downloaded from the NCBI Gene Expression Omnibus (GEO accession GSE98622). Estimated abundance of clusters 0 and 2 were further fitted along time using a generalized additive model to allow nonlinear relationships via thin plate regression splines (MGCV package). The second dataset includes pod mice treated by ABT384 or its vehicle. This dataset is described above. Secondly, we deconvoluted a human bulk-RNAseq dataset. Briefly, 42 allograft kidney recipients were enrolled at the University Hospitals of Leuven. In each case, protocol biopsy was performed at four different time points: before implantation (kidney flushed and stored in ice), after reperfusion (at the end of the surgical procedure) and 3 and 12 months after transplantation^36^. Association between estimated abundance of cluster Fib3 after reperfusion and serum creatinine levels were fitted using a robust linear model. Association between ctci Banff score at one year and estimated abundance of cluster Fib4 at reperfusion was compared using a Wilcoxon test.

### Spatial RNAseq analysis

Six mouse kidneys were analyzed (1 control replicate per genotype; 2 pod28 replicates per genotype) by spatial transcriptomics of 10 μm slices cut on blocks embedded in OCT compounds. After library generation using the Visium Spatial Gene Expression kit (10X Genomics) and sequencing (Functional Genomics Center Zurich, FGCZ). Raw FastQ files were processed using the 10x Genomics Space Ranger counts pipeline, version 2.0.165. This pipeline maps FASTQ reads to a reference genome, filters and counts UMIs against a gene reference, and generates spatially resolved gene expression data. The alignment and counting were performed against the mm10-2020-A mouse reference, consistent with the scRNAseq analysis. To overlay gene expression data onto the corresponding tissue image, high resolution TIFF images of the tissue sections were performed which are essential for the accurate mapping of gene expression data to the spatial coordinates on each tissue. The processed data yielded a matrix of gene counts for each spatial spot, allowing for the investigation of gene expression patterns within the tissue context. For spot clustering analysis and primary annotation, prior to analysis in Seurat, the 10x Genomics Loupe Browser version 7.0 for Visium was utilized for Manual Fiducial Alignment, suppress aberrant dot and preliminary spot annotation based on the ’cortex’, ’medulla’, and ’papilla’ tissue regions. Subsequently, all spot matrices (matrices of gene counts per spot) were imported into R using the Seurat package for graph-based spot clustering analysis and visualization. Spots that were isolated, non-annotated, contained more than 500 or fewer than 7,000 unique detected different transcripts, or had fewer than 1093 1,000 UMI counts, were filtered out. Following this initial filtering, the remaining datasets underwent further processing using the SCTransform function to normalize and stabilize gene expression variance across spots. This step included regressing out the per-spot percentage of mitochondrial transcripts. Canonical Correlation Analysis (CCA) was deployed to project all samples into a shared dimensional space using Seurat. We selected 3,000 integration features to identify anchors across datasets, and employed the SCTransform method for normalization. The first 30 CCA components constituted the neighbors search space, using 20 neighbors for anchor selection. This integration step produced a corrected count matrix for the 3,000 anchor features, which was then used for PCA. The first 30 PCs derived from PCA were utilized for graph-based clustering with the Louvain algorithm. Subsequently, the integrated low-dimensional space was visualized using UMAP, incorporating the first 30 principal components. Spots were annotated and visualized based on their predicted cell-type proportions, following a label transfer process that utilized the refined scRNA-seq annotation as a reference. The proportion of signatures arising from each 55 μm spot. For fibroblast analysis, we determined the proportion of fibroblast signatures corresponding to each of the four experimental conditions (control wt and *Hsd11b1* ko animals; pod28 wt and *Hsd11b1* ko animals) in the ’cortex’ spots of all samples. To examine the localization of fibroblasts, we focused on spots with more than 10% fibroblast signature; these spots are called ’fibroblast’. These spots were then clustered into two groups: spots surrounded only ’fibroblast’ spots, termed ’fibroblast center’ and spots defined as ’fibroblast border’, interpreted as the interface between the fibrosis area and kidney tissue. Within ’fibroblast’ spots, we further selected spots with ADAMTSL1 expression exceeding 0.5, called Fibroblast/Adamtsl1+. In contrast, ’fibroblast’ spots with ADAMSTL1 expression of less than 0.5 are called Fibroblast/Adamtsl1-.

### Oxoreductase activity

11β-HSD1-dependent 11-oxoreductase activity was determined by measuring the conversion of cortisone to cortisol using kidney microsome preparations obtained from C57/BL6 wt and *Hsd11b1* ko animals. Briefly, ¾ of the right kidney was grinded in liquid nitrogen to a fine powder and subsequently homogenized in 1 mL homogenization buffer (20 mM Tris-HCl pH 7.5, 50 mM KCl, 2 mM MgCl_2_, 250 mM sucrose) supplemented with protein inhibitor cocktail (PIC, Roche) in a glass bottle with 20 strokes. Homogenates were centrifuged for 20 min at 4°C and 12,000 g. The supernatant was transferred to Airfuge tubes (Beckmann Coulter) and centrifuged for 1 h at 4°C and 105,000 g. The supernatants were discarded and pellets resuspended in 60 μL resuspension buffer (20 mM MOPS pH 7.2, 100 mM KCl, 20 mM NaCl, 1 mM MgCl_2_, PIC). The BCA Protein Assay Kit (Thermo Fisher Scientific) was used to determine the protein concentration. To measure 11β-HSD1 activity, 0.9 μg/μL protein extract was used from each sample. Resuspension buffer was used for the reaction mix and measured as previously described^78^ with an adjustment for the incubation time to 1 h instead of 10 min.

### Quantification of plasma bile acids

We used the plasma ursodeoxycholyltaurine (UDCA-Tau) to 7-oxolithocholyltaurine (7oxo-LC-Tau) ratio as a biomarker of the efficacy of pharmacological 11β-HSD1 inhibition^79,80^. Briefly, 500μl of retro-orbital capillary blood was collected in lithium/heparin-coated tubes (Sarstedt AG&Co, Germany) and centrifuged for 4 min at 3,500 rpm. About 150 μL of plasma was immediately frozen in liquid nitrogen and subsequently stored at -80°C. UDC-Tau and 7oxo-LC-Tau concentrations were measured as previously described^81,82^. For statistical analysis, parametric data were compared by Student’s t or Mann–Whitney test, depending on their distribution. A P value < 0.05 was considered significant. All P values were two-tailed and adjusted for multiple comparisons using Benjamini–Hochberg correction if necessary.

## Acknowledgments

E.F. received support by the Swiss National Science Foundation (SNSF; grant 310030_212434) . S.N.K. received support by the SNSF (AMBIZIONE grant PZ00P3_179916). A.O. received support by the SNSF (310030_214978). R.H.W. received support by the SNSF (310030_219198). A.P. received support by Innosuisse 44124.1 IP-LS. C.B. received support by the SNSF (310030_182317). D.L. received support by the Geneva University Hospitals (PRD 5-2020-I and PRD 4-2021-II) and G.A. received support by the Fondation Ernst and Lucie Schmidheiny. We warmly thank Claes B. Wollheim (PHYM department, University of Geneva, Switzerland) for critical reading of the manuscript. We thank the Histology and Bioimaging core facilities (PHYM department, University of Geneva, Switzerland) for their precious help.

---

**Extended data Fig. 1.**
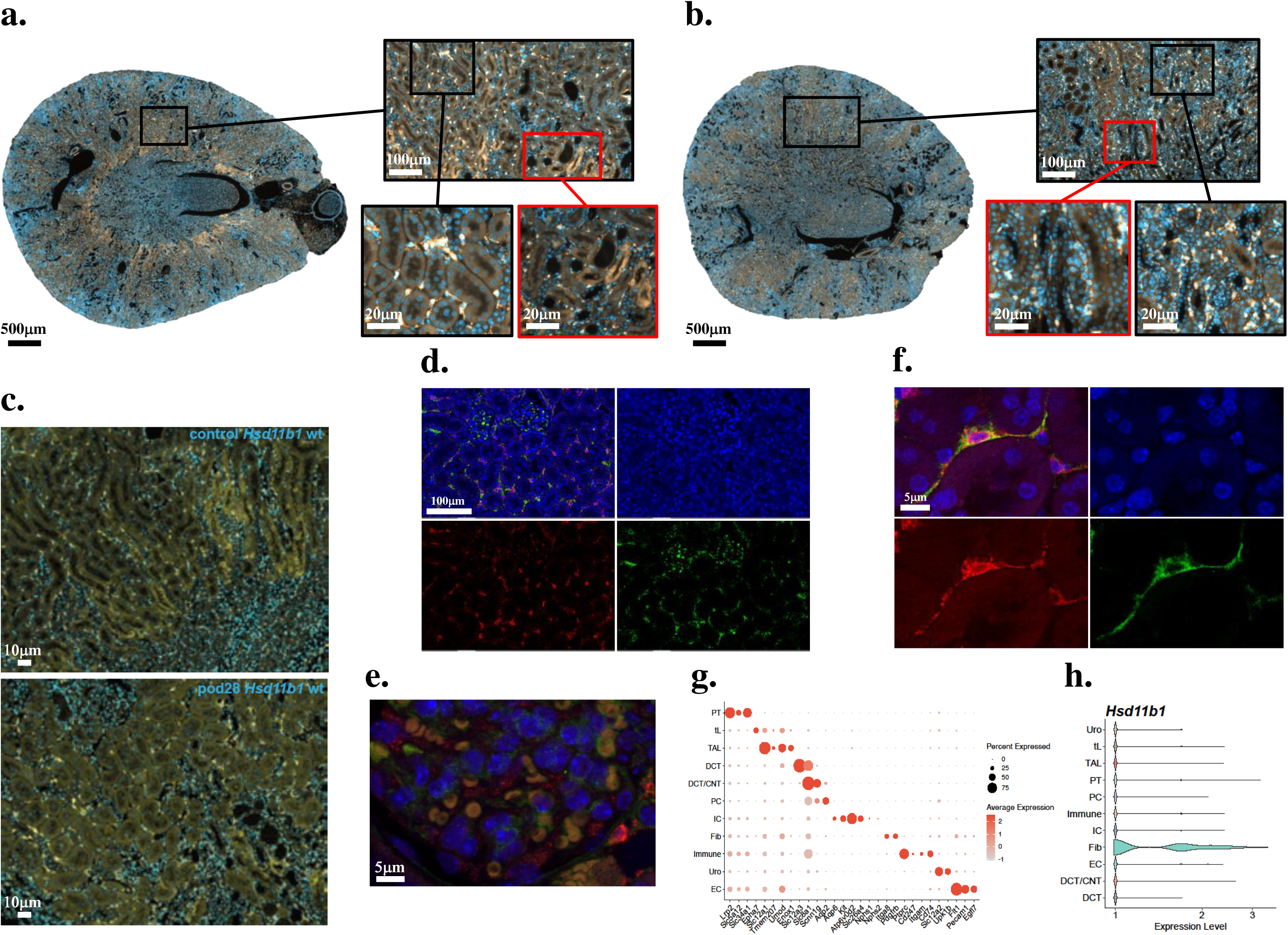
(**a and b**) Representative immunofluorescence images of 11β-HSD1/DAPI in pod mice 7 days post-injury (**a**) and 14 days post-injury (**b**). 11β-HSD1, gold; DAPI, blue. (**c**) Representative cortical confocal images illustrating cortical interstitial cells expressing 11β-HSD1 in control (upper) and pod (lower) mice kidneys. 11β-HSD1, gold; DAPI, blue. (**d to f**) Split 11β-HSD1/PDGFRβ/DAPI channels of representative confocal cortical (**d and e**) and glomerular (**f**) images from non-injured wt mice kidney sections (n=2). 11β-HSD1, red ; PDGFRβ, green ; DAPI, blue. (**g**) Dot plot showing average marker gene expression values and proportion expressed for each cell type in the kidney. (**h**) Violin plots displaying the expression of *Hsd11b1* across identified cell clusters.

**Extended data Fig. 2.**
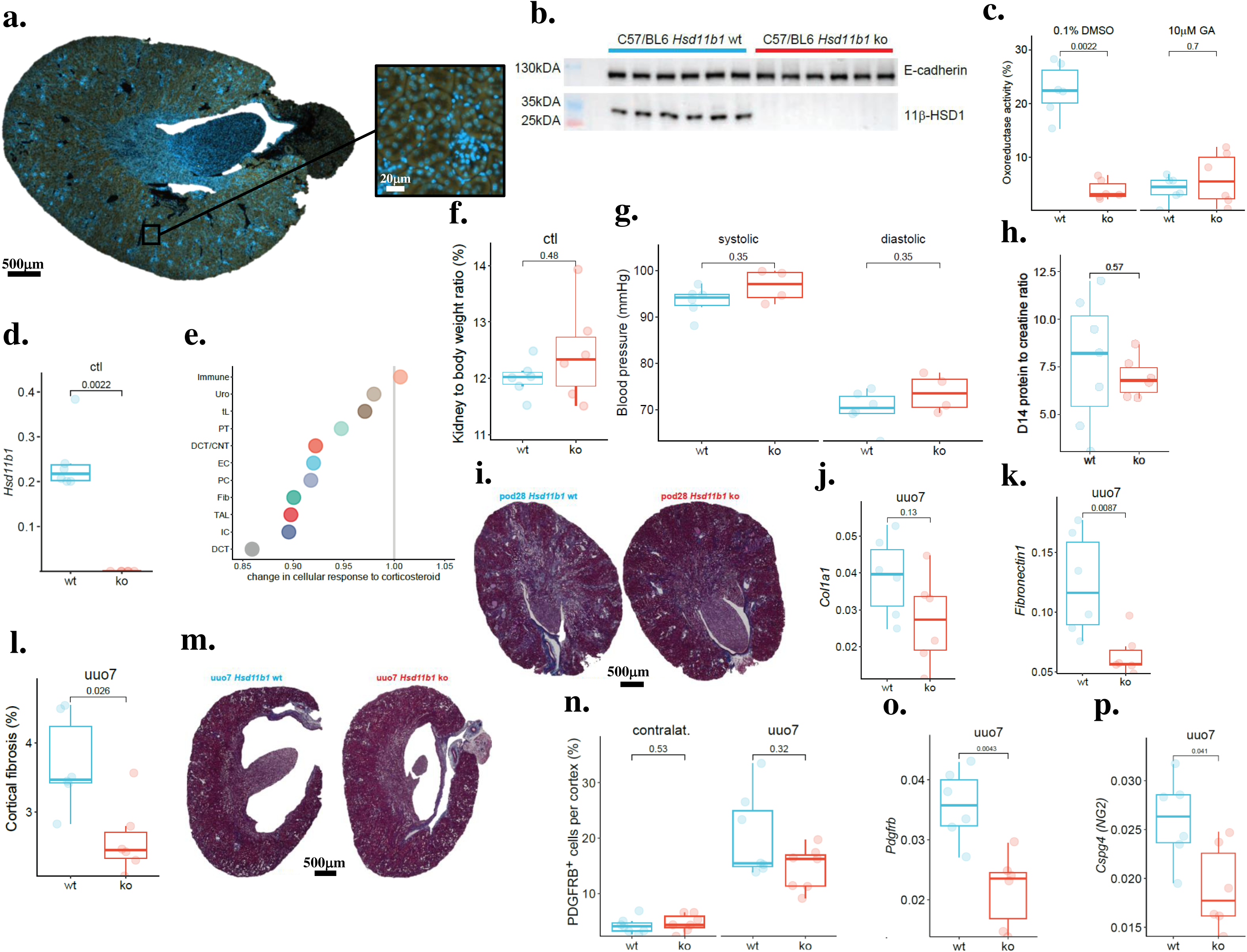
(**a**) Immunofluorescence staining of 11β-HSD1 in non-injured *Hsd11b1* ko kidney sections (n = 1). 11β-HSD1, gold; DAPI, blue. (**b**) Western blot of 11β-HSD1 and E-cadherin in cortical protein extracts of non-injured wt (blue) and ko (red) mice (n = 6). (**c**) Oxoreductase activity measurements (cortisone to cortisol conversion), based on microsome preparations from wt and *Hsd11b1* ko mice kidney tissues with the addition of either 0.1% dimethyl sulfoxide (DMSO) or glycyrrhetinic acid (GA) in 0.1% DMSO (n = 6). (**d**) qPCR analysis of *Hsd11b1* gene expression from wt and ko *Hsd11b1* cortical extracts (n = 6). (**e**) Gene Ontology Biological Process (GOBP) analysis of the cellular response to corticosteroid across clusters from non-injured wt and *Hsd11b1* ko mice. Immune, immune cell; Uro, urothelial cell; tL, thin limb; PT, proximal tubule; DCT/CNT, distal convoluted/connecting tubule; EC, endothelial cell; PC, principal cell; Fib, fibroblast; TAL, thick ascending limb; IC, intercalated cell; DCT, distal convoluted tubule. (**f**) Kidney to body weight ratio (n =6). (**g**) Basal blood pressure measured by tail cuff in non-injured wt and *Hsd11b1* ko mice (n=4-5). (**h**) Measurement of urinary protein at day 14 after injury in wt and *Hsd11b1* ko mice. (**i**) Representative Masson’s trichrome images without overlaid pixel classifiers in pod mice 28 days post-injury. (**j and k**) qPCR analysis of gene expression from wt and *Hsd11b1* ko mice kidney cortical extracts from 7 days obstructed kidneys. Expression levels of *Col1a1* (**j**) and *Fn1* (encoding fibronectin 1) (**k**) genes (n=6). Uuo7, 7 days unilateral ureteral obstruction. (**l and m**) Automated quantification of cortical interstitial fibrosis in obstructed kidneys of wt and *Hsd11b1* ko mice (**l**) and representative Masson’s Trichrome images without overlaid pixel classifiers in 7 days obstructed kidneys (**m**). (**n**) Quantification of interstitial PDGFRβ+ cells in wt and *Hsd11b1* ko 7 days obstructed kidneys and contralateral non-obstructed kidneys (n=6). Contral, contralateral kidney. (**o and p**) qPCR analysis of *Pdgfrb* (**o**) and *Cspg4* (**p**) mRNA levels in cortices of 7 days obstructed wt and *Hsd11b1* ko kidneys (n=6).

**Extended data Fig. 3.**
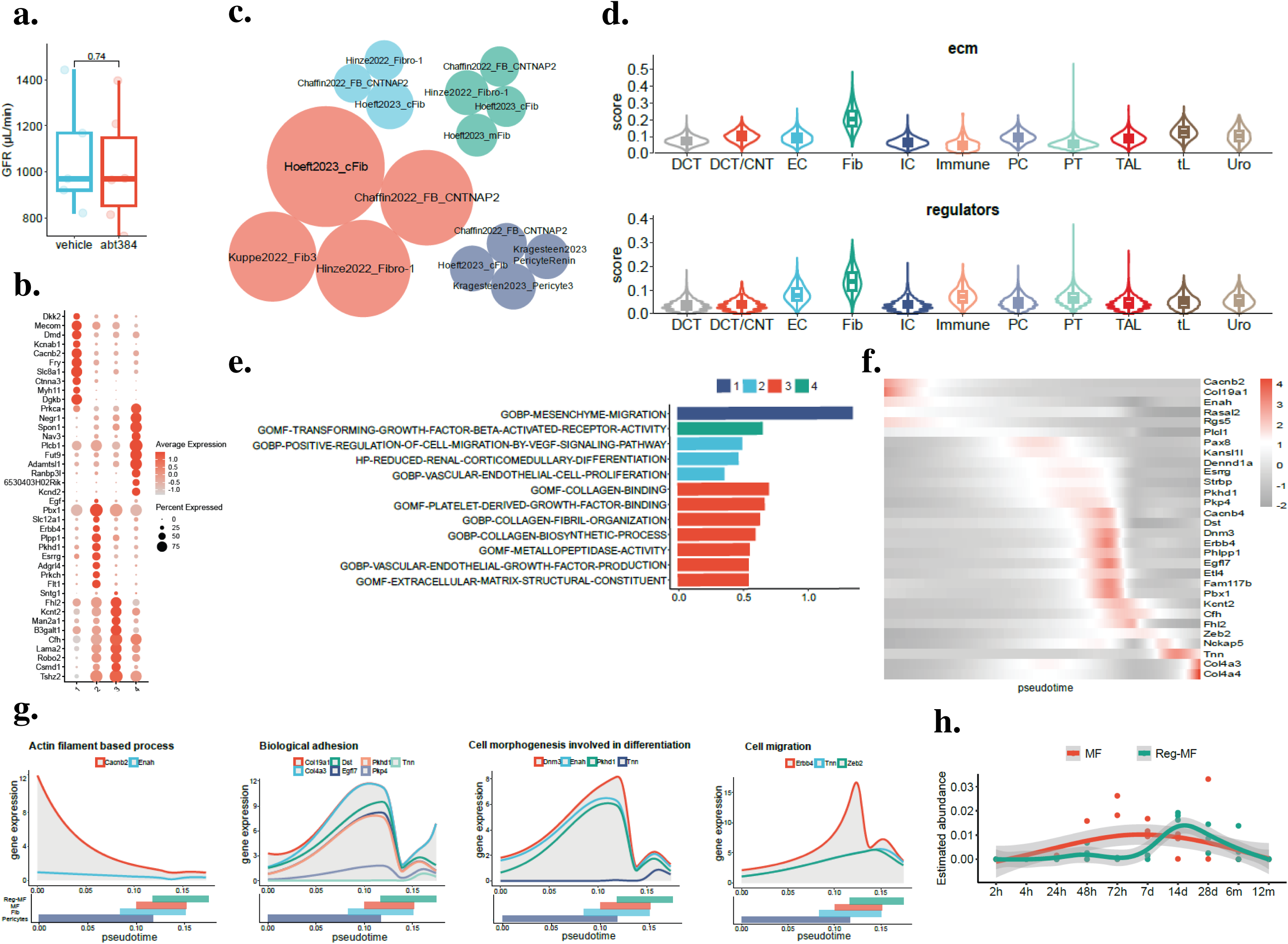
(**a**) Glomerular filtration rate (GFR) (μL/min) at 28 days after induction of glomerular injury. Vehicle, pod28 mice treated with vehicle (0.5% methylcellulose, 0.2% Tween-80) (blue) ; ABT384, pod28 mice treated with vehicle or ABT384 inhibitor (red). (**b**) Dot plot showing the top 10 genes expressed in each of the four mesenchymal clusters identified. *Dkk2*, dickkopf WNT signaling pathway inhibitor 2; *Mecom*, MDS1 and EVI1 complex locus; *Dmd*, dystrophin; *Kcnab1*, potassium voltage-gated channel subfamily A regulatory beta subunit 1; *Cacnb2*, calcium voltage-gated channel auxiliary subunit beta 2; *Fry*, FRY microtubule binding protein; *Slc8a1*, solute carrier family 8 member A1; *Ctnna3*, catenin alpha 3; *Myh11*, myosin heavy chain 11; *Dgkb*, diacylglycerol kinase beta; Prkca, protein kinase C alpha; *Negr1*, neuronal growth regulator 1; *Spon1*, spondin 1; *Nav3*, neuron navigator 3; *Plcb1*, phospholipase C beta 1; *Fut9*, fucosyltransferase 9; *Adamtsl1*, ADAMTS like 1; *Ranbp3l*, RAN binding protein 3 like; *Kcnd2*, potassium voltage-gated channel subfamily D member 2; *Egf*, epidermal growth factor; *Pbx1*, PBX homeobox 1; *Slc12a1*, solute carrier family 12 member 1; *Erbb4*, Erb-B2 receptor tyrosine kinase 4; *Plpp1*, phospholipid phosphatase 1; Pkhd1, PKHD1 ciliary IPT domain containing fibrocystin/polyductin; *Esrrg*, estrogen related receptor gamma; *Adgrl4*, adhesion G protein-coupled receptor L4; *Prkch*, protein kinase C eta; *Flt1*, Fms related receptor tyrosine kinase 1; *Sntg1*, syntrophin gamma 1; *Fhl2*, four and a half LIM domains 2; *Kcnt2*, potassium sodium-activated channel subfamily T member 2; *Man2a1*, mannosidase alpha class 2A member 1; *B3galt1*, beta-1,3-galactosyltransferase 1; *Cfh*, complement factor H; *Lama2*, laminin subunit alpha 2; *Robo2*, roundabout guidance receptor 2; *Csmd1*, CUB and sushi multiple domains 1; *Tshz2*, teashirt zinc finger homeobox 2. (**c**) Transfer learning using previously published mesenchymal signatures displayed by score weight. (**d**) ECM and regulator score levels for all kidney clusters. PT, proximal tubule; tL and TAL, thin and thick ascending limb; DCT, distal convoluted tubule; DCT/CNT, distal convoluted/connecting tubules; PC, principal cell; IC, intercalated cells; Fib, fibroblast; Immune, immune cell; Uro, urothelial cell; EC, endothelial cell. (**e**) Gene Ontology Biological Process (GOBP) analyses for the four mesenchymal clusters. (**f and g**) Expression pattern (**f**) and selected GOBP analyses (**g**) 4 top genes expressed during mesenchymal differentiation. *Cacnb2*, calcium voltage-gated channel auxiliary subunit beta 2; *Col19a1*, collagen type XIX alpha 1 chain; *Enah*, ENAH actin regulator; *Rasal2*, RAS protein activator like 2; *Rgs5*, regulator of G protein signaling 5; *Plcl1*, phospholipase C like 1; *Pax8*, paired box 8; *Kansl1l*, KAT8 regulatory NSL complex subunit 1 like; *Dennd1a*, DENN domain containing 1A; *Esrrg*, estrogen-related receptor gamma; *Strbp*, spermatid perinuclear RNA binding protein; *Pkhd1*, polycystic kidney and hepatic disease 1/polycystin 1; *Pkp4*, plakophilin 4; *Cacnb4*, calcium voltage-gated channel auxiliary subunit beta 4; *Dst*, dystonin; *Dnm3*, dynamin 3; *Erbb4*, Erb-B2 receptor tyrosine kinase 4; *Phlpp1*, PH domain and leucine rich repeat protein phosphatase 1; *Egfl7*, EGF like domain multiple 7; *Etl4*, epilepsy, occipitotemporal lobe and migraine with aura; *Fam117b*, family with sequence similarity 117 member B; *Pbx1*, PBX homeobox 1; *Kcnt2*, potassium sodium-activated channel subfamily T member 2; *Cfh*, complement factor H; *Fhl2*, four and a half LIM domains 2; *Zeb2*, zinc finger E-box binding homeobox 2; *Nckap5*, NCK associated protein 5; *Tnn*, tenascin N; *Col4a3*, collagen type IV alpha 3 chain; *Col4a4*, collagen type IV alpha 4 chain. (**h**) Relative MF and Reg-MF cluster expression pattern analyses in ischemia-reperfusion-injury-related acute kidney injury snRNAseq datasets.

**Extended data Fig. 4.**
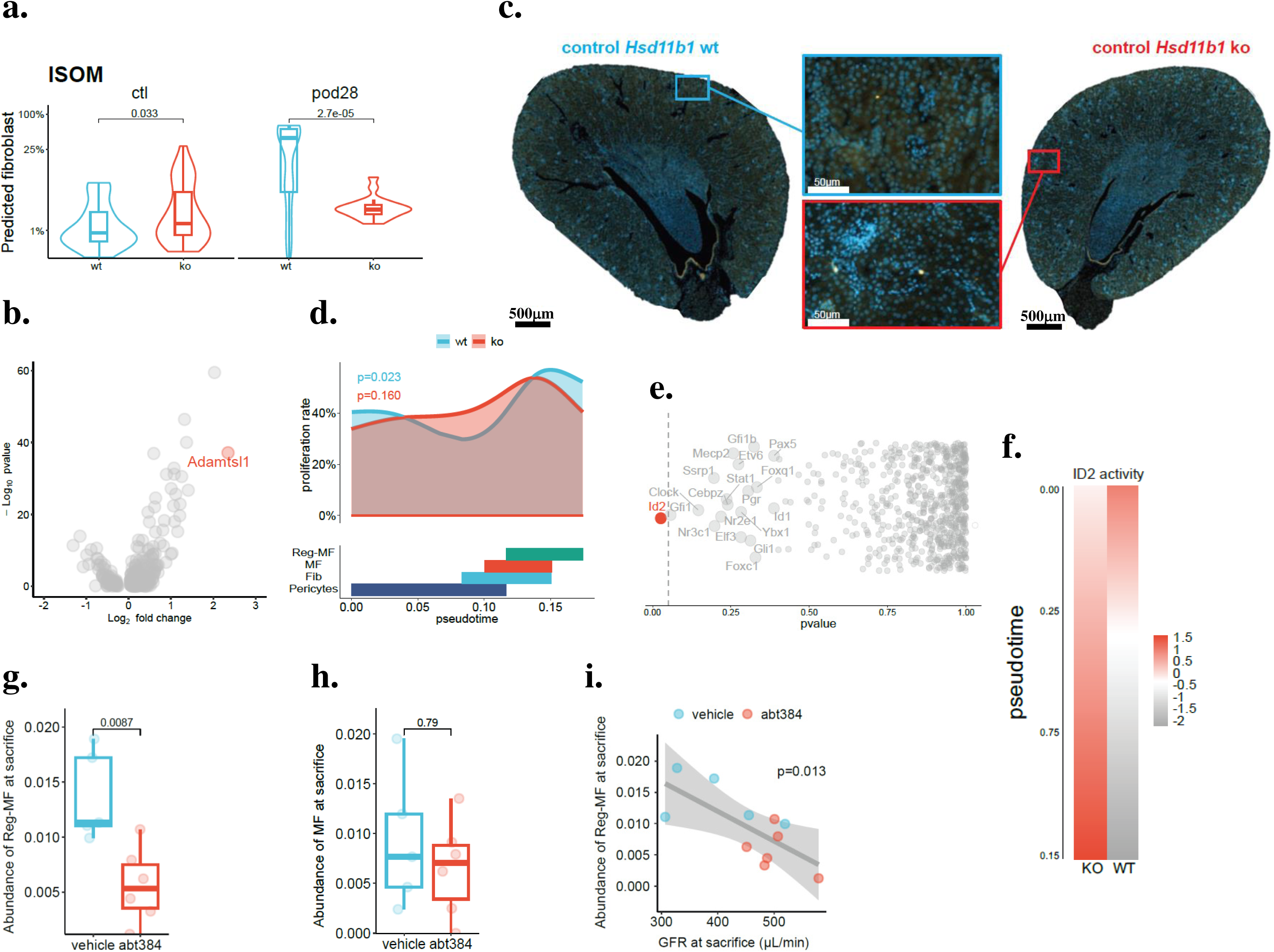
(**a**) Violin plots showing the predicted probability of fibroblast detection in each inner stripe of the outer medulla (ISOM) Visium spot, in control (ctl) and pod 28 days (pod28) animals and among genotypes (*Hsd11b1* wt, blue; *Hsd11b1* ko, red). (**b**) Volcano plot representation showing the most discriminating genes for Reg-MF cluster. (**c**) Representative immunofluorescence staining of ADAMTSL1 in non-injured kidney of wt and *Hsd11b1* ko mice (n = 6). ADAMSTL1, gold; DAPI, blue. (**d**) Proportion of proliferating cells along pseudotime and across genotypes. (**e**) P values for differential transcription factor expression patterns along pseudotime between ko and wt animals. (**f**) Predicted activity of the inhibitor of DNA binding 2 (ID2) transcription factor along pseudotime and across genotypes. (**g**) Abundance of the Reg-MF cluster estimated by bulk RNAseq and deconvolution of snRNAseq mesenchymal cluster signatures. (**h**) Abundance of myofibroblast cluster (MF) estimated by bulk RNAseq and deconvolution of snRNAseq mesenchymal cluster signatures. (**i**) Association between GFR and predicted regulator myofibroblast cluster (Reg-MF) at euthanasia.

**Supplementary table 1.**

| **Variable** | **Beta** | **Pvalue** | **LowCI** | **HighCI** | **Pvalue adj** |
| --- | --- | --- | --- | --- | --- |
| **Donor creatinine** | -2.03E-03 | 7.64E-01 | -1.53E-02 | 1.13E-02 | 9.94E-01 |
| **Donor age** | 1.06E-04 | 3.18E-01 | -1.03E-04 | 3.15E-04 | 9.36E-01 |
| **Donor weight** | 2.07E-04 | 7.85E-02 | -1.62E-05 | 4.30E-04 | 7.44E-01 |
| **Donor length** | 1.21E-06 | 9.94E-01 | -3.13E-04 | 3.16E-04 | 9.94E-01 |
| **Receiver dob** | -1.07E-08 | 9.71E-01 | -5.84E-07 | 5.62E-07 | 9.94E-01 |
| **Receiver age** | -6.53E-06 | 9.51E-01 | -2.15E-04 | 2.01E-04 | 9.94E-01 |
| **Cold ischemia time** | -1.28E-04 | 6.24E-01 | -6.36E-04 | 3.80E-04 | 9.36E-01 |
| **Post-reperfusion time** | 9.61E-02 | 4.96E-01 | -1.82E-01 | 3.74E-01 | 9.36E-01 |
| **Hot ischemia time** | 6.67E-05 | 4.96E-01 | -1.26E-04 | 2.60E-04 | 9.36E-01 |
| **Donor sex male** | -3.68E-04 | 8.92E-01 | -5.63E-03 | 4.89E-03 | 9.94E-01 |
| **Donor type DBD** | -5.23E-03 | 5.63E-01 | -1.52E-02 | 4.77E-03 | 9.36E-01 |
| **Donor type DCD** | -5.29E-03 | 5.63E-01 | -1.59E-02 | 5.33E-03 | 9.36E-01 |
| **Donor hypertension yes** | -6.14E-03 | 1.60E-01 | -1.45E-02 | 2.17E-03 | 8.00E-01 |
| **Receiver sex male** | -5.11E-03 | 9.92E-02 | -1.10E-02 | 8.03E-04 | 7.44E-01 |
| **Baseline immunosuppression** | 2.94E-03 | 3.19E-01 | -2.74E-03 | 8.61E-03 | 9.36E-01 |

**Supplementary table 2.**
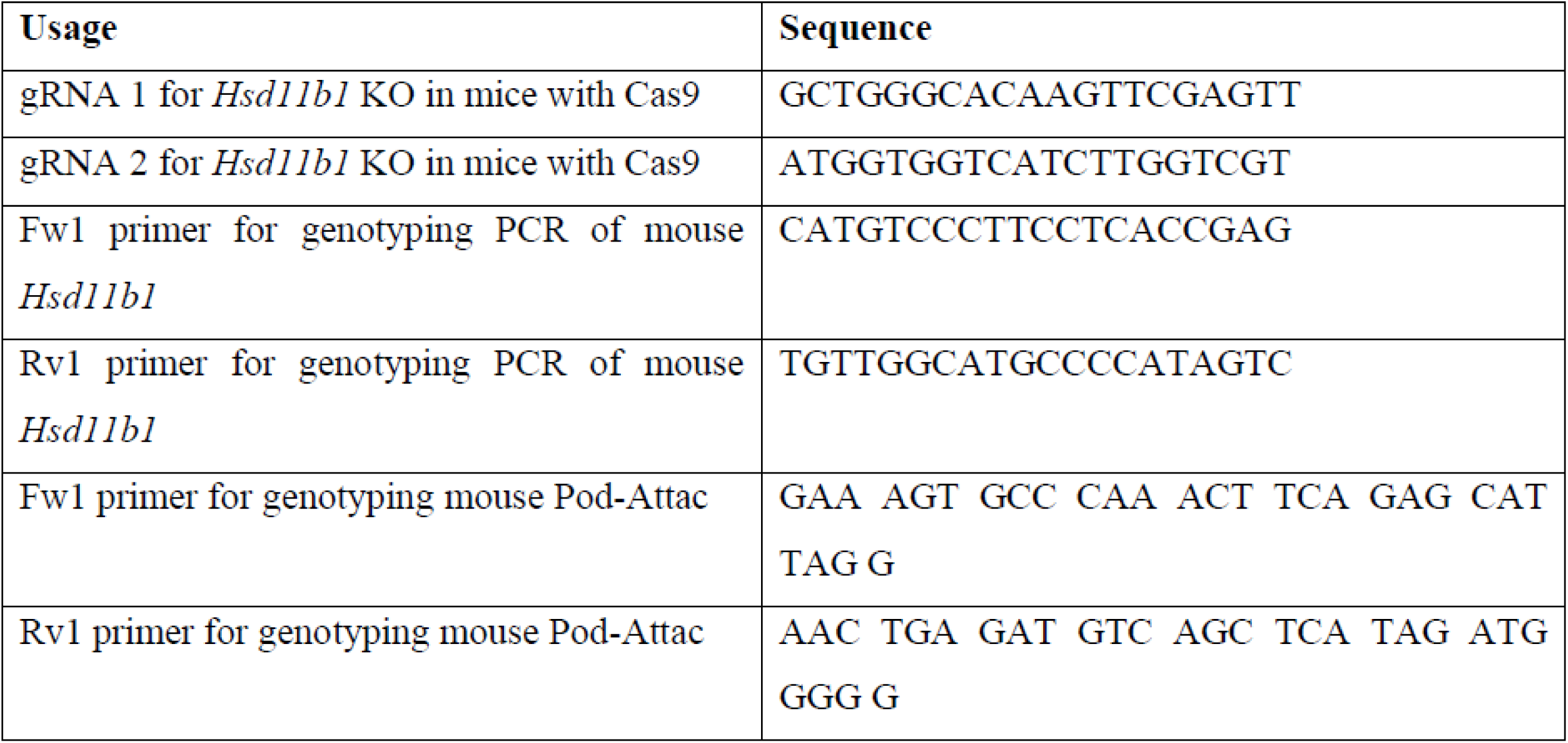

## Bibliography

1. Webster, A. C., Nagler, E. V., Morton, R. L. & Masson, P. Chronic Kidney Disease. Lancet 389, 1238–1252 (2017).

2. Tuttle, K. R. et al. Incidence of Chronic Kidney Disease among Adults with Diabetes, 2015-2020. N Engl J Med 387, 1430–1431 (2022).

3. Li, L., Fu, H. & Liu, Y. The fibrogenic niche in kidney fibrosis: components and mechanisms. Nat Rev Nephrol 18, 545–557 (2022).

4. Hoeft, K. et al. Platelet-instructed SPP1+ macrophages drive myofibroblast activation in fibrosis in a CXCL4-dependent manner. Cell Rep 42, (2023).

5. Risdon, R. A., Sloper, J. C. & De Wardener, H. E. Relationship between renal function and histological changes found in renal-biopsy specimens from patients with persistent glomerular nephritis. Lancet 2, 363–366 (1968).

6. Kuppe, C. et al. Decoding myofibroblast origins in human kidney fibrosis. Nature 589, 281–286 (2021).

7. Kuppe, C. et al. Spatial multi-omic map of human myocardial infarction. Nature 608, 766–777 (2022).

8. Lebleu, V. S. et al. Origin and function of myofibroblasts in kidney fibrosis. Nat Med 19, 1047–1053 (2013).

9. Kramann, R. et al. Perivascular Gli1+ progenitors are key contributors to injury-induced organ fibrosis. Cell Stem Cell 16, 51–66 (2015).

10. Chapman, K., Holmes, M. & Seckl, J. 11β-hydroxysteroid dehydrogenases: intracellular gate-keepers of tissue glucocorticoid action. Physiol Rev 93, 1139–1206 (2013).

11. Odermatt, A. & Kratschmar, D. V. Tissue-specific modulation of mineralocorticoid receptor function by 11β-hydroxysteroid dehydrogenases: an overview. Mol Cell Endocrinol 350, 168–186 (2012).

12. Tiganescu, A. et al. 11β-Hydroxysteroid dehydrogenase blockade prevents age-induced skin structure and function defects. J Clin Invest 123, 3051–3060 (2013).

13. White, C. I. et al. Cardiomyocyte and Vascular Smooth Muscle-Independent 11β-Hydroxysteroid Dehydrogenase 1 Amplifies Infarct Expansion, Hypertrophy, and the Development of Heart Failure After Myocardial Infarction in Male Mice. Endocrinology 157, 346–357 (2016).

14. Faust, H. J. et al. Adipocyte associated glucocorticoid signaling regulates normal fibroblast function which is lost in inflammatory arthritis. Nat Commun 15, 9859 (2024).

15. Gong, R., Morris, D. J. & Brem, A. S. Human renal 11beta-hydroxysteroid dehydrogenase 1 functions and co-localizes with COX-2. Life Sci 82, 631–637 (2008).

16. Buhl, E. M. et al. Dysregulated mesenchymal PDGFR-β drives kidney fibrosis. EMBO Mol Med 12, (2020).

17. Gray, G. A., White, C. I., Castellan, R. F. P., McSweeney, S. J. & Chapman, K. E. Getting to the heart of intracellular glucocorticoid regeneration: 11β-HSD1 in the myocardium. J Mol Endocrinol 58, R1–R13 (2017).

18. Poinot, H. et al. Activation of endogenous glucocorticoids by HSD11B1 inhibits the antitumor immune response in renal cancer. Oncoimmunology 13, (2023).

19. Weingartner, M. et al. The ratio of ursodeoxycholyltaurine to 7-oxolithocholyltaurine serves as a biomarker of decreased 11β-hydroxysteroid dehydrogenase 1 activity in mouse. Br J Pharmacol 178, 3309–3326 (2021).

20. Gómez, C. et al. Identification of a human blood biomarker of pharmacological 11β-hydroxysteroid dehydrogenase 1 inhibition. Br J Pharmacol 181, 698–711 (2024).

21. Kragesteen, B. K. et al. The transcriptional and regulatory identity of erythropoietin producing cells. Nat Med 29, 1191–1200 (2023).

22. Kirita, Y., Wu, H., Uchimura, K., Wilson, P. C. & Humphreys, B. D. Cell profiling of mouse acute kidney injury reveals conserved cellular responses to injury. Proc Natl Acad Sci U S A 117, 15874–15883 (2020).

23. Naba, A. et al. The extracellular matrix: Tools and insights for the ‘omics’ era. Matrix Biol 49, 10–24 (2016).

24. Zhao, H. et al. Secretion of Shh by a Neurovascular Bundle Niche Supports Mesenchymal Stem Cell Homeostasis in the Adult Mouse Incisor. Cell Stem Cell 23, 147 (2018).

25. Kumar, S., et al. ZEB2 controls kidney stromal progenitor differentiation and inhibits abnormal myofibroblast expansion and kidney fibrosis. JCI Insight 8, (2023).

26. Gerhardt, L. M. S. et al. Lineage Tracing and Single-Nucleus Multiomics Reveal Novel Features of Adaptive and Maladaptive Repair after Acute Kidney Injury. J Am Soc Nephrol 34, 554–571 (2023).

27. Duffield, J. S. Cellular and molecular mechanisms in kidney fibrosis. J Clin Invest 124, 2299–2306 (2014).

28. Dimitrov, D. et al. Comparison of methods and resources for cell-cell communication inference from single-cell RNA-Seq data. Nat Commun 13, (2022).

29. Fu, H. et al. Tenascin-C Is a Major Component of the Fibrogenic Niche in Kidney Fibrosis. J Am Soc Nephrol 28, 785–801 (2017).

30. Hao, Y. et al. Dictionary learning for integrative, multimodal and scalable single-cell analysis. Nat Biotechnol 42, 293–304 (2024).

31. Caleb Snider, J., et al. Targeting 5-HT2B Receptor Signaling Prevents Border Zone Expansion and Improves Microstructural Remodeling After Myocardial Infarction. Circulation 143, 1317–1330 (2021).

32. Wang, P. et al. Circular RNA circBNC2 inhibits epithelial cell G2-M arrest to prevent fibrotic maladaptive repair. Nat Commun 13, (2022).

33. Zheng, S. C. et al. Universal prediction of cell-cycle position using transfer learning. Genome Biol 23, (2022).

34. Müller-Dott, S. et al. Expanding the coverage of regulons from high-confidence prior knowledge for accurate estimation of transcription factor activities. Nucleic Acids Res 51, 10934–10949 (2023).

35. Lake, B. B. et al. An atlas of healthy and injured cell states and niches in the human kidney. Nature 619, 585–594 (2023).

36. Cippà, P. E. et al. Transcriptional trajectories of human kidney injury progression. JCI Insight 3, (2018).

37. Breyer, M. D. & Susztak, K. The next generation of therapeutics for chronic kidney disease. Nat Rev Drug Discov 15, 568–588 (2016).

38. Shook, B. A. et al. Myofibroblast proliferation and heterogeneity are supported by macrophages during skin repair. Science 362, (2018).

39. Richeldi, L., Collard, H. R. & Jones, M. G. Idiopathic pulmonary fibrosis. Lancet 389, 1941–1952 (2017).

40. Wiertz, I. A. et al. Unfavourable outcome of glucocorticoid treatment in suspected idiopathic pulmonary fibrosis. Respirology 23, 311–317 (2018).

41. Khan, M. M. et al. Glucocorticoids exacerbate TGFß1 mediated fibrotic signalling in lung fibroblasts by inhibition of metalloproteinase expression and activity. ERJ Open Res 7, 33 (2021).

42. Nakamura, R., Mukudai, S., Bing, R., Garabedian, M. J. & Branski, R. C. Complex fibroblast response to glucocorticoids may underlie variability of clinical efficacy in the vocal folds. Sci Rep 10, (2020).

43. Lavall, D. et al. The mineralocorticoid receptor promotes fibrotic remodeling in atrial fibrillation. J Biol Chem 289, 6656–6668 (2014).

44. Nakamura, T., Girerd, S., Jaisser, F. & Barrera-Chimal, J. Nonepithelial mineralocorticoid receptor activation as a determinant of kidney disease. Kidney Int Suppl (2011) 12, 12– 18 (2022).

45. Barrera-Chimal, J., Bonnard, B. & Jaisser, F. Roles of Mineralocorticoid Receptors in Cardiovascular and Cardiorenal Diseases. Annu Rev Physiol 84, 585–610 (2022).

46. Bridges, J. P. et al. Glucocorticoid regulates mesenchymal cell differentiation required for perinatal lung morphogenesis and function. Am J Physiol Lung Cell Mol Physiol 319, L239–L255 (2020).

47. Zou, X. et al. 11Beta-hydroxysteroid dehydrogenase-1 deficiency or inhibition enhances hepatic myofibroblast activation in murine liver fibrosis. Hepatology 67, 2167–2181 (2018).

48. Xiao, W. et al. 11β-hydroxysteroid dehydrogenase-1 is associated with the activation of hepatic stellate cells in the development of hepatic fibrosis. Mol Med Rep 22, 3191– 3200 (2020).

49. Hirohata, S. et al. Punctin, a novel ADAMTS-like molecule, ADAMTSL-1, in extracellular matrix. J Biol Chem 277, 12182–12189 (2002).

50. Wang, X. et al. Critical Role of ADAMTS2 (A Disintegrin and Metalloproteinase With Thrombospondin Motifs 2) in Cardiac Hypertrophy Induced by Pressure Overload. Hypertension 69, 1060–1069 (2017).

51. Nauffal, V. et al. Genetics of myocardial interstitial fibrosis in the human heart and association with disease. Nat Genet 55, 777–786 (2023).

52. Ajjan, R. A. et al. Oral 11β-HSD1 inhibitor AZD4017 improves wound healing and skin integrity in adults with type 2 diabetes mellitus: a pilot randomized controlled trial. Eur J Endocrinol 186, 441–455 (2022).

53. Othonos, N. et al. 11β-HSD1 inhibition in men mitigates prednisolone-induced adverse effects in a proof-of-concept randomised double-blind placebo-controlled trial. Nat Commun 14, (2023).

54. Rutkowski, J. M. et al. Adiponectin promotes functional recovery after podocyte ablation. J Am Soc Nephrol 24, 268–282 (2013).

55. Khodo, S. N. et al. NADPH-oxidase 4 protects against kidney fibrosis during chronic renal injury. J Am Soc Nephrol 23, 1967–1976 (2012).

56. Dizin, E. et al. Time-course of sodium transport along the nephron in nephrotic syndrome: The role of potassium. FASEB J 34, 2408–2424 (2020).

57. Schreiber, A. et al. Transcutaneous measurement of renal function in conscious mice. Am J Physiol Renal Physiol 303, (2012).

58. Bankhead, P. et al. QuPath: Open source software for digital pathology image analysis. Sci Rep 7, (2017).

59. Ruifrok, A. C., Katz, R. L. & Johnston, D. A. Comparison of quantification of histochemical staining by hue-saturation-intensity (HSI) transformation and color-deconvolution. Appl Immunohistochem Mol Morphol 11, 85–91 (2003).

60. Boothby, I. C. et al. Early-life inflammation primes a T helper 2 cell-fibroblast niche in skin. Nature 599, 667–672 (2021).

61. Hansson, K., Jafari-Mamaghani, M. & Krieger, P. RipleyGUI: software for analyzing spatial patterns in 3D cell distributions. Front Neuroinform 7, (2013).

62. Verissimo, T. et al. PCK1 is a key regulator of metabolic and mitochondrial functions in renal tubular cells. Am J Physiol Renal Physiol 324, F532–F543 (2023).

63. Zheng, G. X. Y. et al. Massively parallel digital transcriptional profiling of single cells. Nat Commun 8, (2017).

64. McGinnis, C. S., Murrow, L. M. & Gartner, Z. J. DoubletFinder: Doublet Detection in Single-Cell RNA Sequencing Data Using Artificial Nearest Neighbors. Cell Syst 8, 329–337.e4 (2019).

65. Young, M. D. & Behjati, S. SoupX removes ambient RNA contamination from droplet-based single-cell RNA sequencing data. Gigascience 9, (2020).

66. Legouis, D. et al. Altered proximal tubular cell glucose metabolism during acute kidney injury is associated with mortality. Nat Metab 2, 732–743 (2020).

67. Wang, L. et al. Single-cell dual-omics reveals the transcriptomic and epigenomic diversity of cardiac non-myocytes. Cardiovasc Res 118, 1548–1563 (2022).

68. Wang, L. et al. Single-cell dual-omics reveals the transcriptomic and epigenomic diversity of cardiac non-myocytes. Cardiovasc Res 118, 1548–1563 (2022).

69. Ianevski, A., Giri, A. K. & Aittokallio, T. Fully-automated and ultra-fast cell-type identification using specific marker combinations from single-cell transcriptomic data. Nat Commun 13, (2022).

70. Bhuva, D. D., Cursons, J. & Davis, M. J. Stable gene expression for normalisation and single-sample scoring. Nucleic Acids Res 48, E113 (2020).

71. Armingol, E. et al. Context-aware deconvolution of cell-cell communication with Tensor-cell2cell. Nat Commun 13, (2022).

72. Baghdassarian, H. M., Dimitrov, D., Armingol, E., Saez-Rodriguez, J. & Lewis, N. E. Combining LIANA and Tensor-cell2cell to decipher cell-cell communication across multiple samples. Cell reports methods 4, (2024).

73. Moon, K. R. et al. Visualizing structure and transitions in high-dimensional biological data. Nat Biotechnol 37, 1482–1492 (2019).

74. Street, K. et al. Slingshot: cell lineage and pseudotime inference for single-cell transcriptomics. BMC Genomics 19, (2018).

75. Hao, Y. et al. Integrated analysis of multimodal single-cell data. Cell 184, 3573–3587.e29 (2021).

76. Wang, X., Park, J., Susztak, K., Zhang, N. R. & Li, M. Bulk tissue cell type deconvolution with multi-subject single-cell expression reference. Nat Commun 10, (2019).

77. Liu, J., et al. Molecular characterization of the transition from acute to chronic kidney injury following ischemia/reperfusion. JCI Insight 2, (2017).

78. Kley, M., Moser, S. O., Winter, D. V. & Odermatt, A. In vitro methods to assess 11β-hydroxysteroid dehydrogenase type 1 activity. Methods Enzymol 689, 121–165 (2023).

79. Weingartner, M. et al. The ratio of ursodeoxycholyltaurine to 7-oxolithocholyltaurine serves as a biomarker of decreased 11β-hydroxysteroid dehydrogenase 1 activity in mouse. Br J Pharmacol 178, 3309–3326 (2021).

80. Gómez, C. et al. Identification of a human blood biomarker of pharmacological 11β-hydroxysteroid dehydrogenase 1 inhibition. Br J Pharmacol 181, 698–711 (2024).

81. Gómez, C., Stücheli, S., Kratschmar, D. V., Bouitbir, J. & Odermatt, A. Development and Validation of a Highly Sensitive LC-MS/MS Method for the Analysis of Bile Acids in Serum, Plasma, and Liver Tissue Samples. Metabolites 10, 1–17 (2020).

82. Weingartner, M. et al. The ratio of ursodeoxycholyltaurine to 7-oxolithocholyltaurine serves as a biomarker of decreased 11β-hydroxysteroid dehydrogenase 1 activity in mouse. Br J Pharmacol 178, 3309–3326 (2021).

